# Heme acts a chloroplast-to-nucleus retrograde signal to regulate intercellular trafficking via plasmodesmata

**DOI:** 10.1101/2025.08.05.668529

**Authors:** Mohammad F. Azim, Levi B. Gifford, Mazem Alazem, Jesse D. Woodson, Tessa M. Burch-Smith

## Abstract

Intercellular communication via plasmodesmata (PD) is essential for plant growth, development, and defense, yet its regulation remains poorly understood. Chloroplasts communicate information about the environment and the physiological state of the plant cell to the nucleus. It has been proposed that chloroplast-generated signals also modulate the expression of nuclear genes related to PD to regulate the trafficking of photosynthetic products and metabolites along with other molecules that act non-cell-autonomously. In this study, we set out to identify the chloroplast retrograde signals that regulate intercellular trafficking via PD. Using a combination of *Arabidopsis thaliana* mutants and gene silencing in *Nicotiana benthamiana,* we found that the metabolites of the tetrapyrrole biosynthetic pathway, most likely heme, can act to modulate PD-mediated intercellular trafficking. We also identified genes that are potentially regulated by the heme signal to modify plasmodesmata function. Together, these findings strengthen the link between chloroplasts and PD in coordinating intercellular communication for optimal plant development and resource allocation.

## INTRODUCTION

Plasmodesmata (PD) are membrane-lined cytoplasmic pores that allow the movement of metabolites and signaling molecules between neighboring plant cells [1–5]. PD connect plant cells to each other to form a symplast, and almost all plant cells are part of the symplast for at least part of their development [6]. In land plants, the central membranous structure of the pore, derived from the endoplasmic reticulum (ER) of the connected cells, is called the desmotubule. The space between the desmotubule and plasma membrane is called the cytoplasmic sleeve or annulus. This cytoplasmic space is viewed as the primary route for the trafficking of soluble molecules between cells [7, 8]. In the last decade or so, much has been learnt about the lipid and protein composition of PD and of their roles as signaling hubs for co-ordination of whole plant responses [9–11]. Despite this compositional complexity, only a simple concept of plasmodesmal function in intercellular trafficking persists.

Despite being integral to the plant cellulosic cell wall, plasmodesmal frequency is not a fixed cellular parameter, and the number of PD in a cell wall changes during cell wall expansion and in response to developmental and environmental signals, presumably to maintain a necessary minimal level of intercellular communication [3, 12, 13]. This is exemplified by PD converting from simple to branched forms during the sink-to-source transition in leaves, where sink leaves possess predominantly simple PD while source leaves PD adopt complex configurations that often include multiple pores connected by a central cavity, the so-called branched PD [14]. The cellular signals and pathways that direct these changes in PD remain poorly understood. The organelle-nucleus-PD signaling (ONPS) hypothesis attempts to explain at least some of the regulation of PD form and function in response to developmental and other signals. According to ONPS, signals originating in the mitochondria and chloroplast, acting alone or together, regulate the expression of cell wall- and PD-related nuclear genes to determine plasmodesmal structure and permeability, ultimately controlling PD-mediated trafficking [12, 15]. Since chloroplasts are the sites of photosynthesis carbon fixation, it is reasonable to postulate that chloroplasts would regulate PD-mediated trafficking to control the allocation of fixed carbon between source and sink tissues. Studies of the chloroplast RNA helicase ISE2 underpin the ONPS hypothesis and connect chloroplast gene expression with regulation of PD trafficking. Knockout of *ISE2* in *Arabidopsis thaliana* mutants or knockdown of *ISE2* expression in *Nicotiana benthamiana* increased intercellular trafficking via PD and stimulated the biogenesis of new PD [4, 12, 16]. ISE2 has multiple roles in plastid RNA processing including promoting splicing of group II introns, post-transcriptional cytidine to uridine (C-to-U) RNA editing, and ribosomal RNA processing and accumulation [17, 18]. In keeping with ISE2’s role in plastid gene expression, Arabidopsis *ise2* mutants or *N. benthamiana* leaves where *ISE2* was silenced exhibited defective chloroplast biogenesis and development [12]. Indeed, it seems that any defects in chloroplast gene expression or other processes disrupting chloroplast biogenesis and development can result in altered PD development and intercellular trafficking, supporting the ONPS hypothesis [18–21]. However, the signaling mechanisms employed by ISE2 and chloroplasts for ONPS remain unresolved.

Chloroplast biogenesis from proplastids or etioplasts requires massive changes in the expression of nuclear genes. The suite of genes whose expression correlates with chloroplast development includes not only photosynthesis-associated nuclear genes (PhANGs) [22] but also genes directing cell wall synthesis and/or modification, sugar transport, stress responses, and responses to reactive oxygen species (ROS) [23]. Signals from chloroplasts to the nucleus during chloroplast biogenesis that regulate nuclear gene expression are called biogenic chloroplast retrograde signals (CRS) [24]. Biogenic CRS pathways have been identified by inhibiting normal chloroplast development. For instance, when chloroplast development is disturbed by herbicides (e.g., norflurazon) or by blocking plastid translation (e.g., lincomycin), the nucleus responds by reducing the expression of PhANGs like *LIGHT HARVESTING CHLOROPHYLL A/B BINDING PROTEIN1.2 (LHCB1.2)* [25, 26]. This seminal observation was the basis of forward genetic screens that identified mutants which fail to repress expression of PhANGs, revealing a breakdown in coordination between the chloroplast and nuclear gene expression. The mutants were consequently named the *<u>g</u>enomes <u>un</u>coupled* (*gun*) mutants. The original genetic screens for mutants with defects in chloroplast-to-nucleus retrograde signaling identified six loci (*gun1-6*) that are important in retrograde signaling. Five of these (*gun2-6*) encode components of the chloroplast-localized tetrapyrrole biosynthesis (TBS) pathway [27–29], while the sixth, *gun1*, encodes a chloroplast-localized pentatricopeptide repeat (PPR) protein [30]. *GUN2*, *GUN3*, and *GUN5* encode heme oxygenase, phytochromobilin synthase, and the H subunit of Mg-chelatase (CHLH), respectively [29], while *GUN4* encodes a regulator of Mg-chelatase activity [28].

The TBS pathway produces critical molecules for the production the iron-containing cofactor, heme, and the magnesium (Mg) containing chlorophylls [31]. The branchpoint of the TBS pathway – where protoporphyrin (ProtoIX) can be a substrate for either Mg-chelatase or Fe-chelatase - is highly regulated to prevent the accumulation of the photosensitizing and singlet oxygen-producing intermediates [32]. It was originally proposed that Mg-protoporphyrin IX (Mg-proto IX) accumulated in undeveloped chloroplasts and acts as a CRS to negatively regulate nuclear gene expression [33]. This hypothesis was challenged by additional biochemical and genetic analyses that demonstrated that bulk Mg-Proto levels did not correlate with PhANG expression [34, 35]. Instead, the identification of the dominant *gun6* mutation (*gun6-D*) that resulted in elevated FERROCHELATASE (FC)1 activity led to the hypothesis that the FC1 product, heme, was a positive regulator of PhANGs [27]. Heme has been established as a signaling molecule in other systems like algae [36], yeast [37], *C. elegans* [38]*, and humans* [39]. In the case of algae, it can be exported from chloroplasts [40–42].

The role of the TBS pathway and its intermediates in influencing PD-mediated intercellular trafficking was examined in this study. In *A. thaliana gun* mutants and in *N. benthamiana* plants where expression of *GUN* and heme-related genes was silenced, we found that changes in the TBS pathway flux was correlated with intercellular trafficking. Indeed, the data suggest that total heme levels, and not other TBS intermediates, are important for altering plasmodesmal function. Gene expression analyses in this system allowed us to identify a suite of nuclear genes that are potentially responsive to heme to modify plasmodesmal function. Supporting this observation, changes in intercellular trafficking were positively correlated with plasmodesmal density and not with callose metabolism at PD. These results provide further support for the role of heme, a highly conserved electron-carrier across all living systems, as an important signaling molecule. Heme has been proposed to have critical roles beyond oxygen binding and electron transfer in several biological systems [36–39]. The proposed connection between the heme/the TBS pathway and PD-mediated intercellular trafficking reported in this work highlights the deep evolutionary connections between PD and chloroplasts [15, 43, 44]. Further understanding of how tetrapyrroles control trafficking through PD could enable the engineering of plants with optimized carbon allocation for higher yield or carbon partitioning to limit pathogen spread.

## RESULTS

### GUN5 has a role in regulating intercellular trafficking in Arabidopsis and *N. benthamiana*

Our previous study found that silencing genes encoding proteins with roles in chloroplast gene expression and biogenesis resulted in defective PD-mediated intercellular trafficking as well as altered expression of several genes of the TBS pathway including *GUN2-5* [20]. Given that the TBS pathway provides retrograde signals for chloroplast biogenesis [27], we hypothesized that this pathway influences intercellular trafficking. Indeed, GUN2 and GUN3 belong to the Fe branch of the TBS pathway, while GUN4 and GUN5 are part of the Mg branch [45] **(Fig 1A).** To investigate whether disruption of the TBS pathway impacts PD-mediated trafficking a GFP movement assay was used to measure intercellular trafficking in Arabidopsis *gun* mutants. For this, a plasmid for *GFP* expression was introduced into leaves by low-pressure particle bombardment and the size of foci containing GFP was measured 24 hours later. The extent of intercellular movement of GFP from the primary transformed cells into neighboring cells was expressed as the number of surrounding rings of cells, i.e., Foci of given ring size or mean size of GFP foci **(Fig. 1B** and **1C)**. While *gun2* and *gun4* mutants did not exhibit significant changes in GFP movement, *gun5* mutants had decreased PD-mediated intercellular trafficking revealed as a 20% reduction in mean focus size, from ∼ 2 cell layers in the non-silenced controls to ∼1.4 cell layers in *gun5* mutants (**Fig 1B**).

**Fig 1.**
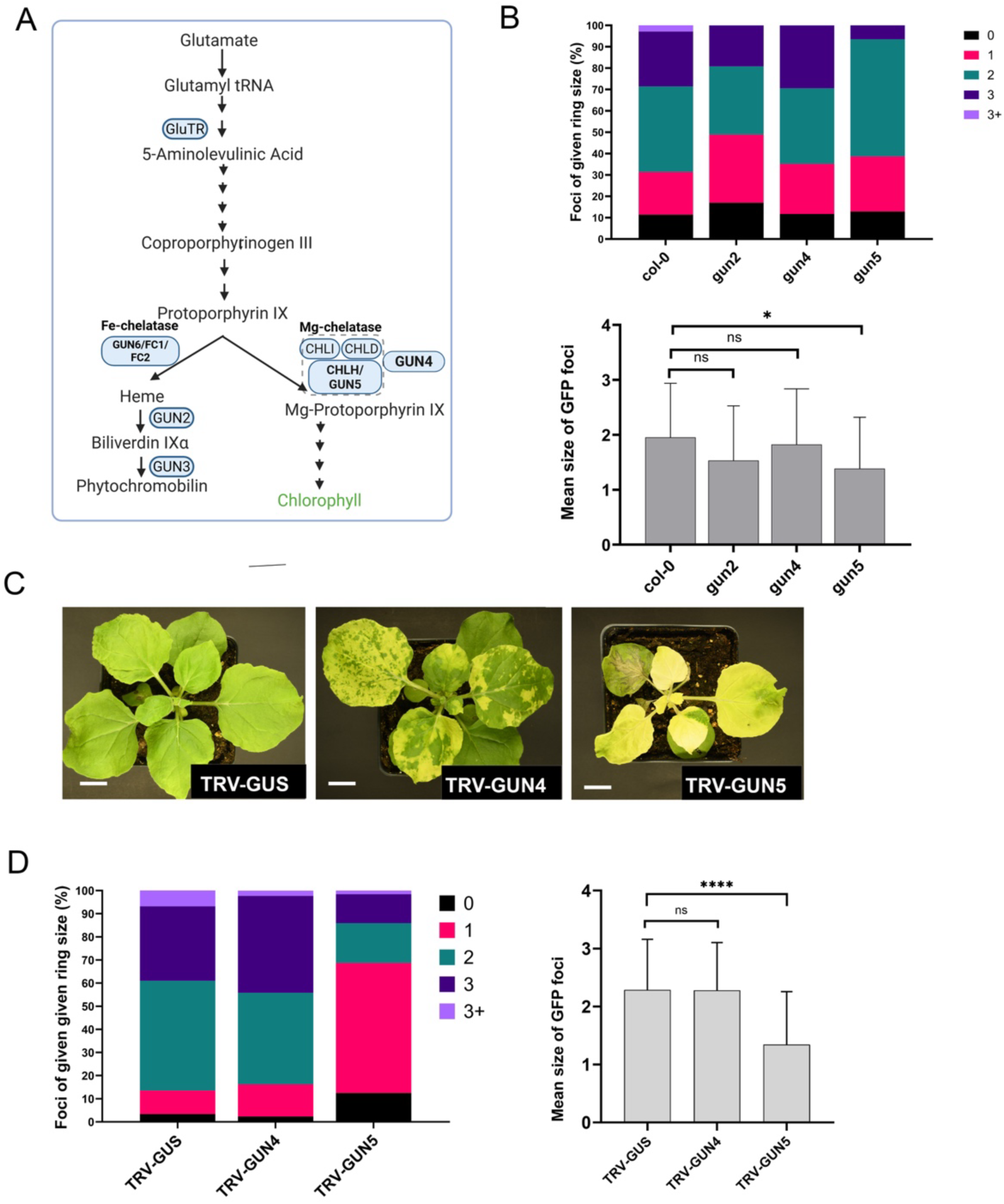
*GENOME UNCOUPLED (GUN)* genes have distinct roles in PD-mediated trafficking in *N. benthamiana*. **(A)** Illustration of the TBS pathway featuring *GUN-*encoding enzymes. The four Mg-chelatase subunits (CHLI, CHLD, CHLH/GUN5, and GUN4) are highlighted. **(B)** PD-mediated trafficking was measured using a movement assay tracking GFP’s diffusion from a single cell into adjacent cells (called rings) after particle bombardment. Stacked bar graph presents the percentage of foci of different ring sizes (0,1,2,3 or >3). The bar graph presents the mean ring size. N=3, n=35. Statistical significance compared to non-silenced control (TRV-*GUS)* was calculated by Mann-Whitney U test. *=p<0.05. (C) VIGS of *GUN4 (*TRV-*GUN4*) caused a variegated phenotype while VIGS of *GUN5 (*TRV-*GUN5*) resulted in severe chlorosis in *N. benthamiana* plants. **(D)** GFP movement assay in silenced plants was performed by Agroinfiltration. Stacked bar graph presents the percentage of foci of different ring sizes and bar graph presents the mean ring size. Statistical significance compared to non-silenced control (TRV-*GUS)* was calculated by Mann-Whitney U-test with significance at p<0.0001 (indicated by **** asterisks). N=4, n=60. Error bars indicate +/- standard deviation.

Since GUN4 and GUN5 are part of the Mg chelatase complex but appeared to have distinct effects on PD-mediated trafficking in the *gun* mutants, their roles in trafficking were more closely examined. For this, *GUN4* and *GUN5* were separately silenced in *Nicotiana benthamiana* by tobacco rattle virus (TRV)-mediated virus-induced gene silencing (VIGS). The *N. benthamiana* genome encodes two homoeologs each for *GUN4* and *GUN5* (**S1 and S2 Fig**), so the VIGS constructs were designed to silence both homoeologs of each gene. Silencing of *GUN4* and *GUN5* was confirmed by quantitative reverse transcription followed by PCR (qRT-PCR) and expression of the of target genes was reduced by as much as 90% compared to expression in a non-silenced control plant (TRV-*GUS*) (**S3 Fig**). The reduced levels of chlorophyll *a*, chlorophyll *b*, and carotenoids measured in the leaves of silenced plants (**S4 Fig**) were consistent with the observed leaf chlorosis, which ranged from a variegated phenotype in *GUN4-*silenced plants to severe yellowing in *GUN5*-silenced plants (**Fig 1C**). GFP movement assays using Agrobacterium to introduce plasmids for *GFP* expression into leaves revealed that silencing *GUN4* did not perturb intercellular trafficking, while *GUN5*-silenced plants had a drastic and statistically significant decrease in PD-mediated trafficking (**Fig 1D**). Thus, GUN4 and GUN5 appear to have distinct roles in regulating intercellular trafficking even though they participate in the same step of the TBS pathway (**Fig. 1A**). Further, these results suggest that GUN5 positively regulates intercellular trafficking.

### ISE2’s involvement in PD-mediated trafficking may require a via GUN5-dependent signaling pathway

ISE2 has important roles in chloroplast RNA metabolism including splicing, C-to-U editing and ribosomal RNA processing is associated with increased PD-mediated intercellular trafficking [16–18, 46]. Given that ISE2, GUN4, and GUN5 have distinct roles in chloroplast development, we investigated whether ISE2 and GUN5 exert their effects on PD via a common signaling pathway. For this, we co-silenced *ISE2* with *GUN4* (TRV-*ISE2*-*GUN4*) and *ISE2* with *GUN5* (TRV-*ISE2*-*GUN5*). Leaves of *ISE2*-silenced plants showed mild chlorosis as expected from previous experiments [12, 20] (**Fig 2A**). When *ISE2* was co-silenced with *GUN4* (TRV-*ISE2*-*GUN4*), leaves were mildly chlorotic like those of *ISE2*-silenced plants and variegation was greatly reduced. Slightly reduced levels of chlorophyll a, chlorophyll b, and carotenoid content (**S4 Fig)** support mild chlorosis in these silenced tissues. In contrast, severe chlorosis, similar to that of plants silenced for *GUN5* alone (**Fig 1C**), was observed when *ISE2* was co-silenced with *GUN5* in TRV-*ISE2*-*GUN5*-silenced plants (**Fig 2A**). Furthermore, when *ISE2* was co-silenced with both *GUN4* and *GUN5* (TRV-*ISE2-GUN4-GUN5*), severe chlorosis was observed, like the *GUN5*-silenced or *ISE2-GUN5*-silenced plants (**Fig 2A**). The very low levels of chlorophyll *a* and *b*, and carotenoids in these plants were consistent with severe chlorosis (**S4 Fig).** Despite their severe chlorosis, all the silenced plants grew and matured to set seed.

**Fig 2.**
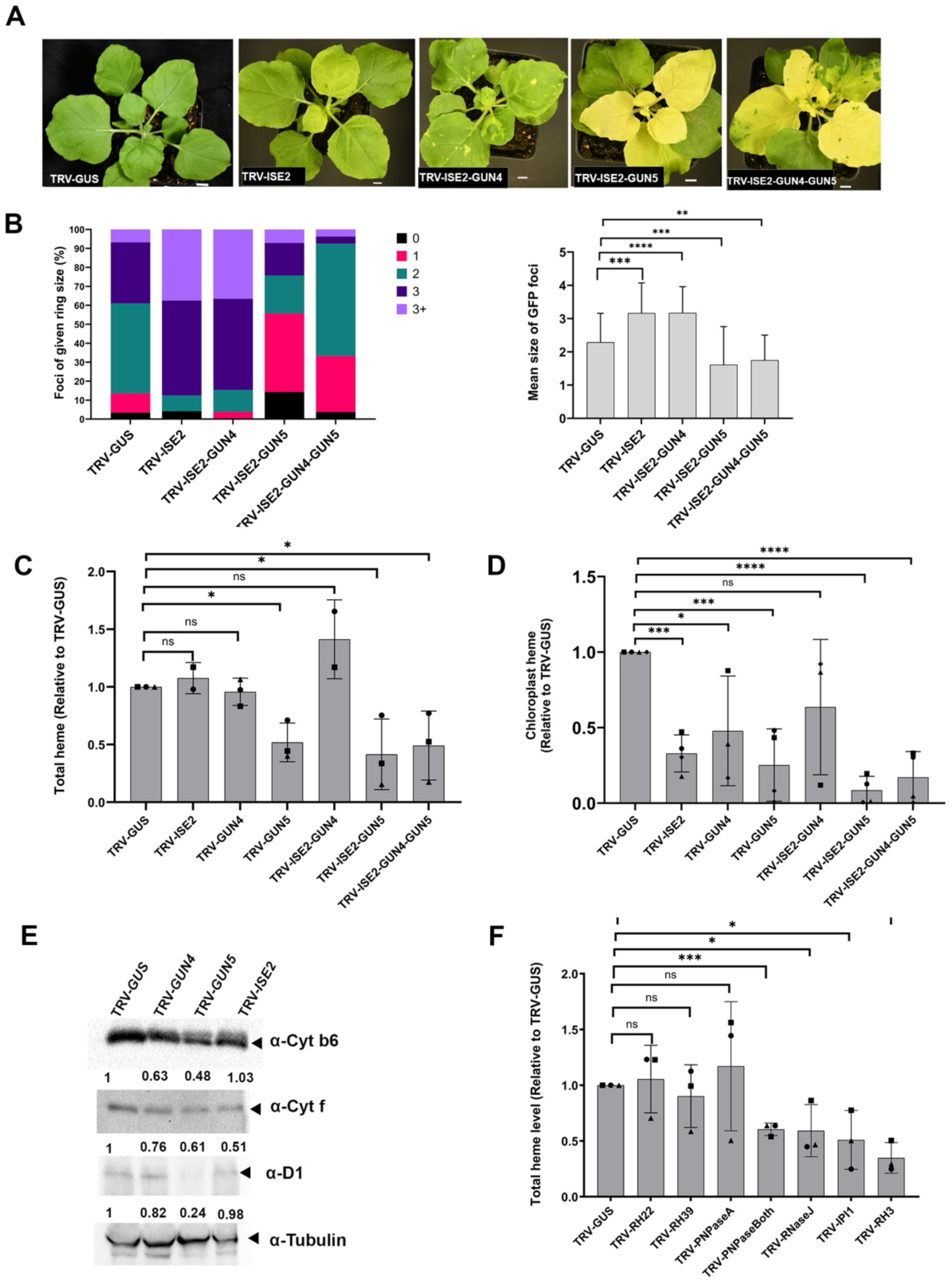
PD-mediated intercellular trafficking is affected by heme levels in *N. benthamiana*. Silencing *ISE2-* only (TRV-*ISE2*) or with *GUN4-* (TRV-*ISE2-GUN4*) and *GUN5-* (TRV-*ISE2-GUN5*) results in different levels of chlorosis.(A) *ISE2-* and *ISE2-GUN4-* silenced plants have mild chlorosis, while *GUN5-* and *ISE2-GUN5-* and *ISE2-GUN4-GUN5-* (TRV-*ISE2-GUN4-GUN5*) silenced plants have sever chlorosis. (B-C) GFP movement in these plants are shown in stacked bar charts showing all data (B) and in (C) as mean layer of GFP movement. Statistical significance compared to the non-silenced control (TRV-*GUS)* was performed by the Mann-Whitney U-test. The steady-state level of non-covalently bound total heme (E) are measured from *N. benthamiana* in non-silenced control (WT-TRV) versus TRV-*ISE2,* TRV-*GUN4,* TRV-*GUN5,* TRV- *ISE2-GUN4,* TRV-*ISE2-GUN5, and* TRV-*ISE2-GUN4-GUN5.* (E) Total heme levels were measured per mg of fresh weight (FW) and are shown relative to non-silenced control (WT-TRV). (F) Total heme levels were also measured after silencing previously known genes (TRV-*RH22,* TRV-*RH39,* TRV-*ISE2,* TRV-*PNPaseA,* TRV-*PNPaseBoth* (silenced both homologs of *PNPases: PNPaseA* and *PNPaseB)*, TRV-*RNaseJ,* TRV- *IPI1,* TRV-*RH3)* that regulate PD-mediated trafficking. Three biological replicates (each replicate contains tissue pooled from three plants) were used to quantify total heme and protoporphyrin IX. Student’s t-test was performed to determine the statistical significance (p<0.05).

To test if retrograde signaling is initiated in the silenced plants, we measured the expression of the photosynthesis-associated nuclear genes *LHCB1.2* and *LHCB2.1* encoding the antennae proteins of the photosystem-associated light-harvesting complexes. Expression of *LHCB1.*2 and *LHCB2.1* was suppressed in *ISE2*-silenced plants (**S5 Fig**). This suggests that disrupting chloroplast RNA processing by silencing *ISE2* triggered retrograde signals. Silencing *GUN4* significantly reduced the expression of only *LHCB2.1* but it did not affect the expression of *LHCB1.2* (**S5 Fig**). In contrast, silencing *GUN5* reduced the expression of both *LHCB1.2* and *LHCB2.1* (**S5 Fig**). Co-silencing *ISE2* with *GUN4* (TRV-*ISE2*-*GUN4*-), with *GUN5* (TRV-*ISE2*-*GUN5*-), and with *GUN4*-*GUN5*- (TRV-*ISE2*-*GUN4*-*GUN5*-) suppressed the expression of *LHCB1.2* and *LHCB2.1* (**S5 Fig**). These results suggest that co-silencing of *ISE2* and/or *GUN5* significantly disrupted chloroplast development and affected CRS to downregulate PhANGs.

GFP movement assays were performed to measure the effects of silencing various combinations of genes on intercellular trafficking. As expected, in *ISE2*-silenced leaves, the GFP movement increased compared to the non-silenced control (TRV-*GUS*), indicating increased PD-mediated trafficking (**Fig 2B**). When *ISE2* was co-silenced with *GUN4* in TRV*-ISE2-GUN4* plants, increased trafficking was observed (**Fig 2B**), which suggests that *ISE2* is epistatic to *GUN4*. In contrast, co-silencing of *ISE2* with *GUN5* in TRV*-ISE2-GUN5* plants resulted in decreased trafficking (**Fig 2B**), indicating that *GUN5* is epistatic to *ISE2*. Taking the phenotypic and trafficking results together, it seems that *ISE2* is epistatic to *GUN4* while *GUN5* is epistatic to *ISE2*. This relationship is supported by the intercellular trafficking measured in plants where *ISE2* was co-silenced with both *GUN4* and *GUN5* (TRV-*ISE2-GUN4-GUN5*). In those plants, trafficking was similar to that measured in *GUN5* or *ISE2*/*GUN5*-silenced plants (**Fig 2B**). Together, these results suggest that ISE2 regulates PD-mediated trafficking via a GUN5-dependent signaling pathway.

### Heme levels are reduced in plants with decreased intercellular trafficking

In plant cells, heme is bound covalently to c-type cytochromes, and non-covalently to b-type cytochromes and most other hemoproteins, including cytochromes P450, nitrate reductase, NADPH oxidases, peroxidases, and catalases [47]. It is generally assumed that there is a pool of free heme that may be involved in signaling, and this pool is likely to be small compared with total cellular heme. It has been proposed that a heme signal may act in the nucleus directly or via interaction with cytoplasmic signaling [48]. We measured the total heme content of leaves from plants where *ISE2* alone or *ISE2* combined with *GUN4* and/or *GUN5* were silenced. Total heme levels in *ISE2*, *GUN4,* and *ISE2-GUN4-*silenced plants were not significantly reduced compared to those in the non-silenced control (TRV-*GUS)* plants (**Fig 2C**). In contrast, leaves from *GUN5-, ISE2-GUN5,* and *ISE2-GUN4-GUN5-*silenced plants contained significantly lower total heme amounts than non-silenced control plants (**Fig 2C).** The reduced heme levels in plants with reduced *GUN5* expression is consistent with previous findings in *N. tabacum* [49]. These findings point to a correlation between reduced levels of total heme and decreased PD-mediated trafficking.

To examine whether a specific pool of heme plays a role in regulating PD-mediated trafficking, we measured heme extracted from isolated chloroplasts, and the heme level was normalized to the total number of chloroplasts (**Fig 2D**). Silencing *ISE2*, *GUN5* or *ISE2-GUN5* reduced chloroplast heme levels compared to the non-silenced control (TRV-*GUS*) plants **(Fig 2D).** Notably, *GUN4-silenced* plants also had a significant reduction in chloroplast heme levels. Thus, the levels of total heme, and not the heme content of the chloroplasts, seem to be important for determining intercellular trafficking capacity, and plants with total heme below a certain threshold seem to reduce their intercellular trafficking.

The effects of silencing *GUN4* and *GUN5* were further characterized by measuring effects on chloroplast proteins (**Fig 2E**). Levels of the heme-associated proteins Cytf and Cytb6 were reduced in plants with reduced *GUN5* expression supporting changes in heme allocation in those silenced plants. There was also a marked reduction in the amount of the photosynthetic center protein, D1, in samples from the *GUN5*-silenced plants, suggesting global defects in the photosynthetic apparatus. The limited effects of silencing *GUN4* on both leaf phenotype and intercellular trafficking were supported by modest changes in protein levels. In samples from *ISE2*-silenced leaves the levels of the Cytf protein were drastically reduced but levels of the Cytb6 protein were unchanged suggesting that changes in protein levels were likely not due to heme deficiency but rather production of specific proteins.

To independently test the apparent correlation between total heme and PD-mediated trafficking, we turned to plants in which chloroplast RNA metabolism genes which were shown to regulate PD-mediated trafficking were silenced. We measured the total heme level after silencing these genes, including *RH3, RH22, RH39, IPI1, RnaseJ,* and two homologs of *PNPases* (*PNPaseA* and *PNPaseB)* as was done previously [20]. The total heme levels of leaves from *RH22-, RH39-, ISE2-, and PNPaseA*-silenced plants previously reported to increase PD-mediated trafficking were not different from those of non-silenced control plants (TRV-*GUS*) **(Fig 2F).** In contrast, leaves from plants where both *PNPases, RnaseJ, IPI1, or RH3* were silenced had reduced levels of total heme **(Fig 2F**). These plants with decreased total heme content have reduced intercellular trafficking [20] like that observed in *GUN5*-silenced plants (**Fig 1B, 1D** and **2B**). Taken together, these findings support that reduced total heme level is correlated with decreased PD-mediated trafficking. They also suggest that a certain minimum level of heme is necessary to be able to increase intercellular trafficking.

### Other TBS pathway intermediates do not correlate with changes in intercellular trafficking

The involvement of other tetrapyrrole intermediates in addition to or besides heme in regulating PD-mediated trafficking was also investigated. We quantified the coproporphyrin III (copro-III) and protoporphyrin IX (proto-IX) levels in leaves of plants where *GUN4* and *GUN5* were individually silenced or co-silenced with *ISE2*. Proto IX is at the branching point, where it can either be converted into heme via ferrochelatase or into other tetrapyrroles in the Mg-branch, while copro-III is upstream of the branching point. Measuring proto-IX and copro-III will help determine whether changes in heme levels are due to branch point or upstream metabolic regulation. The proto IX content was reduced in *ISE2-, GUN5-,* and *ISE2-GUN5*-silenced plants but did not change significantly in other silenced plants **(S6A Fig).** *GUN5*-silenced and *ISE2-GUN5*-silenced plants contained reduced levels of copro-III compared to the non-silenced control (TRV-*GUS)* plants while the copro-III remained unchanged in other silenced plants **(S7A Fig)**. Thus, no correlation between the amount of these TBS pathway intermediates and intercellular trafficking was apparent.

### Distinct pools of heme uniquely affect PD-mediated trafficking

A specific pool of heme generated from the TBS pathway acts as a positive signal that promotes the expression of genes required for chloroplast development [27]. The chloroplast-localized enzymes ferrochelatase 1 (FC1) and FC2 are hypothesized to produce two functionally distinct pools of heme [42]. The bulk of FC2-synthesized heme appears to be utilized mostly within the chloroplast and associates with pETC proteins. FC1 synthesized heme appears to be mostly used for functions outside the chloroplast [42] and is the pool hypothesized to regulate *PhANG* expression during chloroplast development [27]. We therefore set out to determine which pool of heme contributed the signal that modulated PD-mediated intercellular trafficking. Like with the GUN proteins, *FC1* was represented by two homoeologs in the *N. benthamiana* genome (**S8 Fig**). VIGS constructs to silence both homoeologs and one to silence the single copy of *FC2* were designed and *FC1* and *FC2* were silenced individually with the goal of disrupting the two different pools of heme. *FC1*-silenced plants showed no visible phenotype and were indistinguishable from non-silenced control plants (**Fig 3A**). The total heme content as well as the chloroplast heme content were both reduced in the *FC1*-silenced plants, although the decrease in total heme was small (**Fig 3B** and **3C**). Intercellular trafficking was not affected in *FC1*-silenced compared to non-silenced controls (**Fig. 3D**). Like the other TBS pathway genes examined in this work, *FC2* was present as two homoeologs in the *N. benthamiana* genome (**S9 Fig**). In contrast, *FC2-*silenced plants showed severe chlorosis and cell death in the youngest silenced leaves (**Fig. 3A and S10 Fig**). The necrotic lesions on the *FC2*-silenced leaves increased in size over leaf development resulting in necrotic patches when the leaves were fully expanded **(S10 Fig)**. The phenotype of *FC2*-silenced plants is consistent with Arabidopsis *fc2* mutants [50, 51] [42] and tobacco plants where *FC* expression was silenced [52]. Leaves of *FC2*-silenced plants contained significantly reduced levels of both chloroplast and total heme, with a relatively large reduction in total heme compared to the *FC1*-silenced plants (**Fig 3C** and **3D).** However, in contrast to *FC1*-silenced leaves, PD-mediated intercellular trafficking was significantly decreased in *FC2*-silenced leaves (Fig 3D). *GUN2* encodes HEME OXYGENASE that degrades heme [29], and silencing *GUN2* may lead to accumulation of heme [42]. *GUN2*-silenced plants showed mild chlorosis (**Fig. 3A)**, and total heme levels were similar to those of non-silenced control leaves (**Fig 3B**) although chloroplast heme content was significantly decreased (Fig 3C), possibly due to feedback inhibition of ALA biosynthesis (**Fig. 1A**) [53]. PD-mediated intercellular trafficking in *GUN2*-silenced plants was unchanged from that of non-silenced control leaves (**Fig 3D**). Consistent with findings from Espinas et al., [42], levels of the heme-associated proteins Cytb6 and Cytf were reduced in the *FC2*-silenced plants compared to the non-silenced control plants (**Fig 3E**). The levels of these proteins were unaffected in the *FC1-* or *GUN2-*silenced plants. The amount of the photosynthetic center protein, D1, was also drastically reduced in the FC2-silenced plants, indicating broad effects on the proteins in photosynthesis.

**Fig 3.**
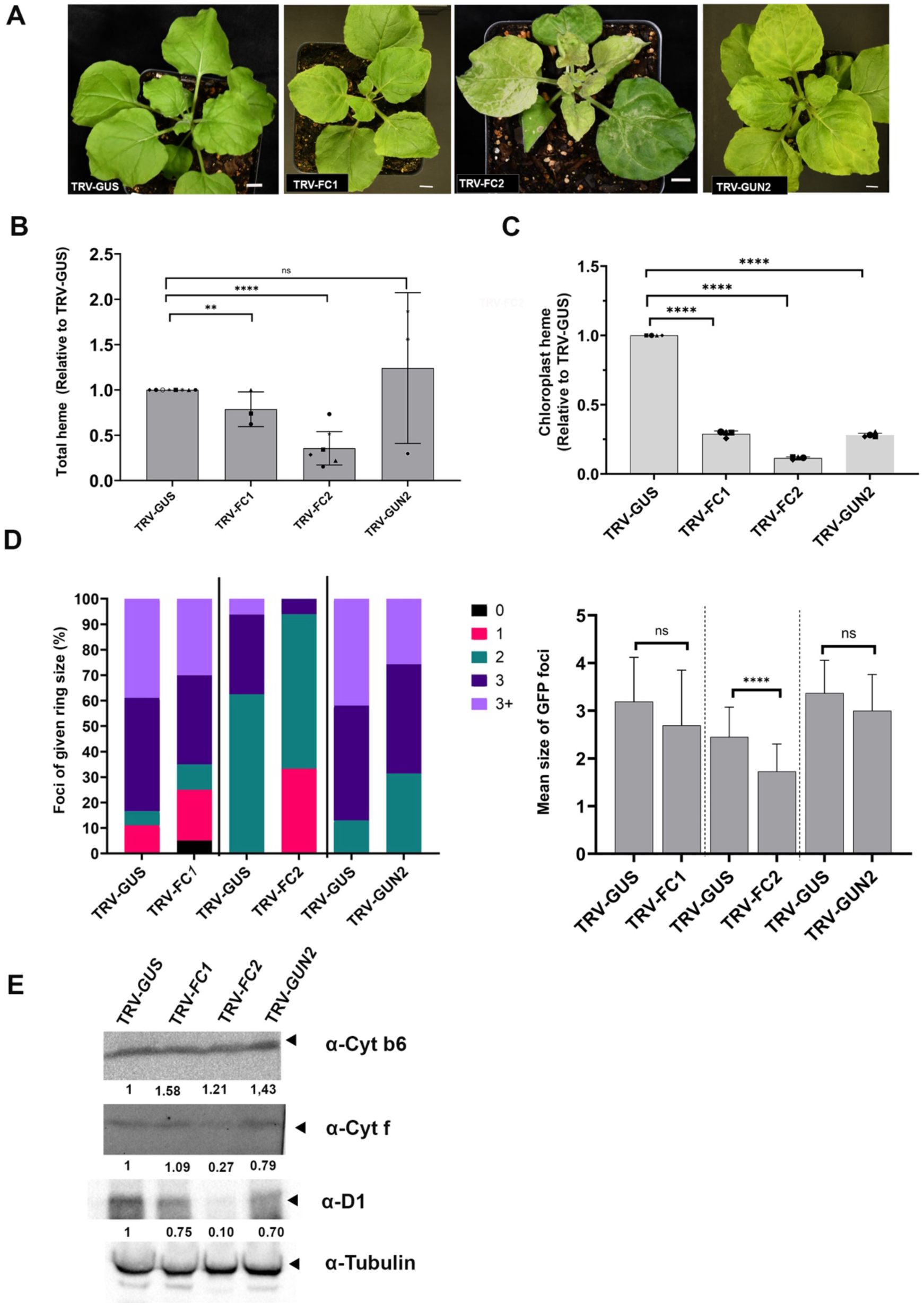
Heme metabolism genes have distinct role in changing intercellular trafficking through PD in *N. benthamiana*. (A) Silencing heme biosynthetic genes *FC1* (TRV-*FC1*) and *FC2-* (TRV-*FC2*) and heme-degrading enzyme encoding gene *GUN2* (TRV-*GUN2*) results in different levels of chlorosis. (A) *FC1-*silenced plants exhibit no chlorosis, while *FC2-*silenced plants show severe chlorosis and *GUN2-*silenced plants show mild chlorosis. (B) Total heme level was measured from a pool of leaves from silenced plants. Three biological replicates (each replicate contains tissue pooled from three plants) were used to quantify total heme. (C) Chloroplast heme was measured from isolated chloroplasts and normalized by the number of the chloroplasts. Total heme (B) and chloroplast heme (F) were measured per mg of fresh weight (FW) and are shown as relative to non-silenced control (WT-TRV). Statistical analyses were performed by Student’s t-test (p <0.05). (D) GFP movement is shown in stacked bar charts presenting all data (left) and the same data as mean area of GFP movement (right). Statistical significance compared to the non-silenced control (TRV-*GUS)* was determined by the Mann-Whitney U-test.

We also measured the proto-IX and copro-III content in *FC1-, FC2-*, and *GUN2*-silenced plants and found no statistically significant difference in any silenced plants compared to the non-silenced controls (**S6B and S7B Fig)**. Together, these results support the finding that heme, and not other TBS intermediates (copro-III and proto-IX), may signal to control intercellular trafficking via PD. Further, they suggest that a heme pool produced by FC2 is critical for maintaining PD-mediated intercellular trafficking.

### Application of exogenous heme increased the PD-mediated trafficking

Our data suggested that heme may act in signaling to regulate intercellular trafficking. The endogenous level of heme is tightly controlled through feedback inhibition of the synthesis of 5-aminolevulinic acid (ALA), a key precursor in the TBS pathway (**Fig. 1**). This feedback loop creates a bottleneck that limits heme accumulation. Indeed, silencing *GUN2* failed to cause accumulation of heme in silenced plants (**Fig. 3B** and **C**). To investigate whether heme was sufficient to regulate intercellular trafficking and to bypass the endogenous block to heme accumulation, we infiltrated plants with heme over a range of concentrations (0, 5, 25, 50, and 100 µM) and performed GFP movement assays in the infiltrated leaves. We hypothesized that if heme could indeed regulate intercellular trafficking, this approach would increase intercellular trafficking by increasing the heme levels in the treated plant. A low concentration of heme (5 µM) increased intercellular trafficking compared to the mock treatment (KOH only) (**Fig. 4A).** Higher heme concentrations of 25 µM and 50 µM resulted in even larger increases in intercellular trafficking compared to the mock treatment (**Fig. 4A)**. The highest concentration of heme tested, 100 µM, did not change intercellular trafficking compared to the mock-treated control (**Fig. 4A**), possibly due to the negative cellular effects of high levels of heme including lipid oxidation [54]. The observed effects of heme on intercellular trafficking agree with the measurement of lower total heme levels in plants with reduced intercellular trafficking. The simplest interpretation of these observations is that, at least in *N. benthamiana*, the infiltration of heme increased cellular heme levels and this led to increased intercellular trafficking.

**Fig 4.**
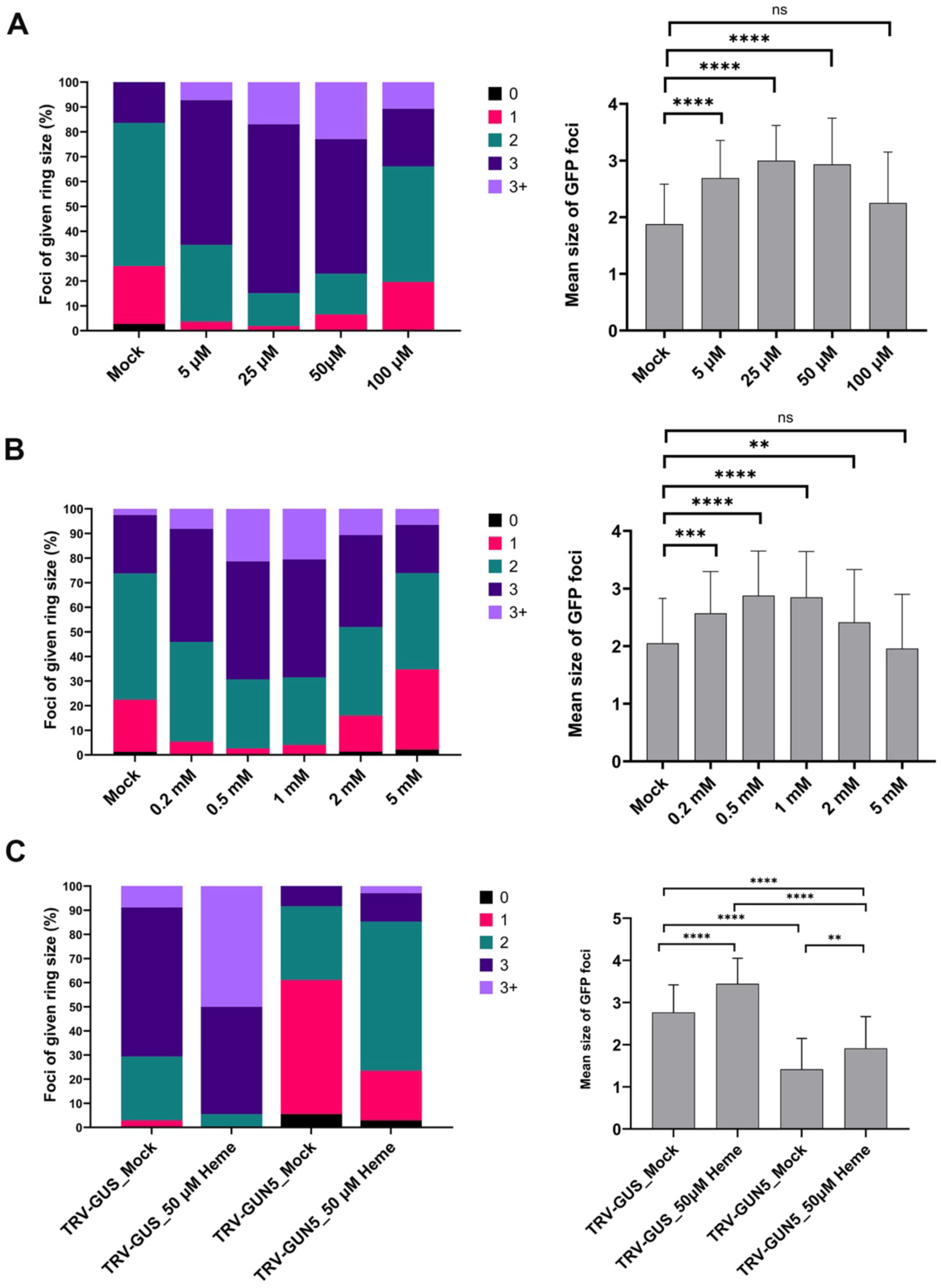
Exogenous application of heme and ALA increase intercellular trafficking in *N. benthamiana*. (A) Intercellular trafficking was measured after infiltrating with 5 µM to 100 µM hemin. GFP movement is shown in stacked bar charts presenting all data (left) and the same data as mean area of GFP movement (right). Statistical significance compared to the non-silenced control (TRV-*GUS)* was determined by the Mann-Whitney U-test. (B) Application of the ALA also caused a significant increase in intercellular trafficking over a range of concentrations from 0.2 to 2 mM, while 5 mM ALA infiltration did not increase trafficking statistically significantly. Movement data presented as described for (A). (C) Infiltration of 50 µM of heme, not 1 mM ALA, partially rescued PD-mediated trafficking in *GUN5-*silenced plants. Movement data presented as described for (A). Statistical significance was measured by the Mann-Whitney U-test. * p<0.05, **p<0.005, ***p<0.0005 (asterisks). ns = p>0.05. Error bars indicate +/- standard deviation.

In a second strategy to overcome the bottleneck of heme accumulation due to the negative feedback to ALA, we treated plants with ALA. This approach was demonstrated to bypass feedback inhibition and promote increased heme biosynthesis [55]. We infiltrated different concentrations (0, 0.2, 0.5, 1, 2, and 5 mM) of ALA into plants and then measured intercellular trafficking in ALA-treated leaves. At a concentration of ALA of 0.2 mM, intercellular trafficking was significantly increased compared to the mock treatment (phosphate buffer) (**Fig. 4B**). Increasing ALA concentration to 0.5, 1 or 2 mM significantly increased intercellular trafficking (**Fig. 4B**). In contrast, 5 mM ALA, had no effect on intercellular trafficking compared to the mock-treated control. The observed increased trafficking with increased ALA concentrations supports the observation that increased heme levels, up to a point, can increase intercellular trafficking.

Finally, we predicted that application of heme could rescue the intercellular trafficking defect in silenced plants with reduced heme levels. To test this, we infiltrated *GUN5*-silenced plants with heme or ALA. Exogenous application of heme at 50 µM increased intracellular trafficking compared to the mock treatment (KOH only) in *GUN5-*silenced plants (**Fig. 4C**). However, it did not completely restore intercellular trafficking to the levels observed in non-silenced plants. Taken together, these experiments suggest that heme can act as the CRS to regulate intercellular trafficking via PD.

### Heme regulates PD-mediated trafficking in a light-independent manner

To examine the interplay between light and the TBS intermediates in PD-mediated signaling we treated leaves with heme then exposed them to standard light (120 μmol m^-2^ · s^−1^) or kept them in the dark before measuring GFP movement. In non-silenced control leaves (TRV-*GUS*), exogenous heme significantly increased trafficking through PD in leaves kept under both light or dark conditions, suggesting that light is not required for heme’s function in PD-mediated trafficking (**Fig 5A**, compare mock_Heme_Light vs 50mM heme_light and mock_Heme_Dark vs 50mM heme_Dark). Similarly, in *GUN5*-silenced plants exogenous heme significantly increased trafficking through PD in leaves kept under light or in the dark, suggesting heme’s function in these *GUN5*-silenced plants is independent of light (**Fig 5B**, compare mock_Heme_Light vs 50mM heme_light and mock_Heme_Dark vs 50mM heme_Dark).

**Fig 5.**
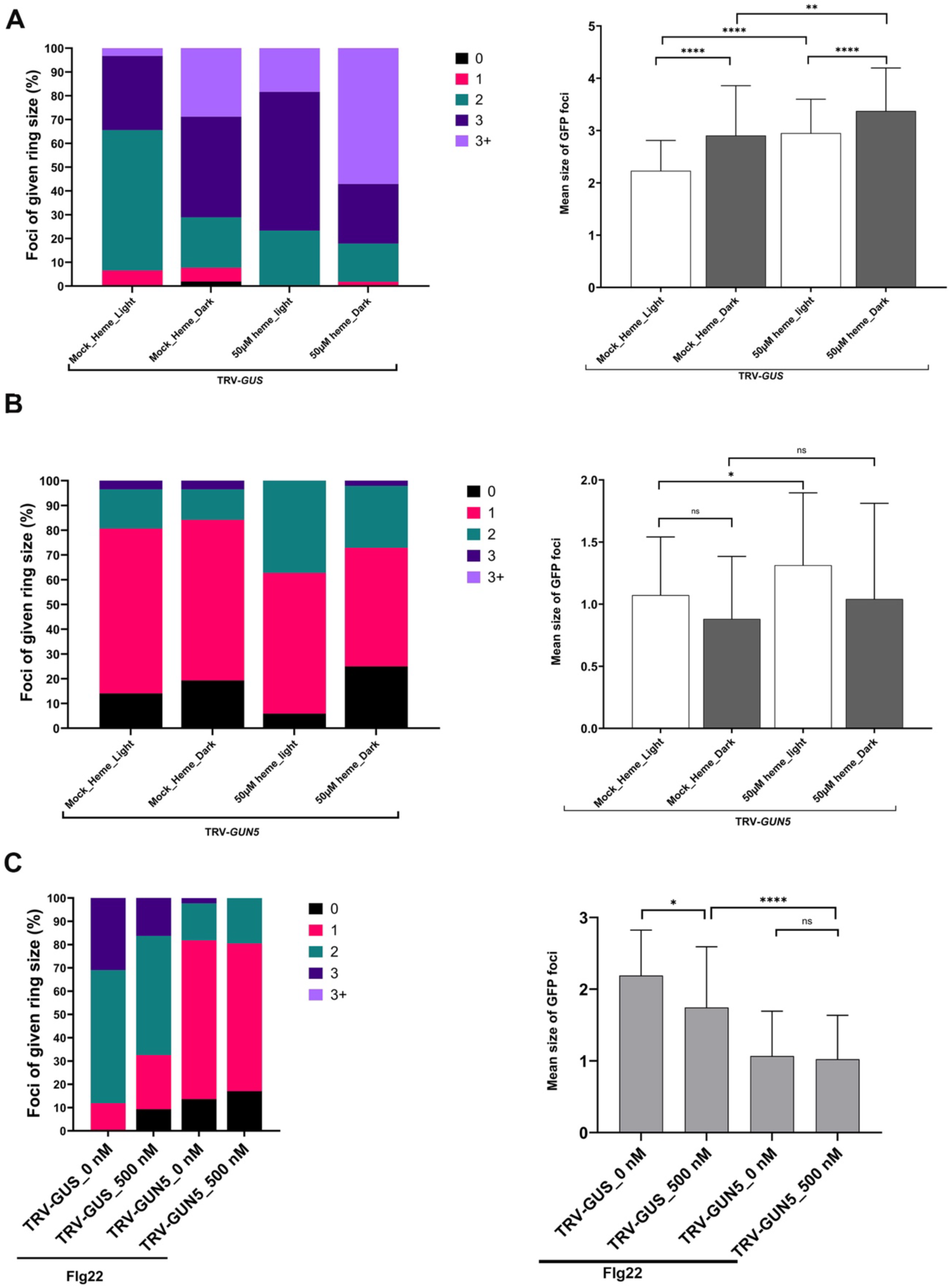
Light dependence of exogenous heme in regulating PD-mediated trafficking in *N. benthamiana*. (A) GFP movement assays for non-silenced control (TRV-*GUS*) plants. Stacked bar charts showing all data (left) and as mean area of GFP movement (right). Mock (0 µM heme, buffer only) or 50 µM heme were infiltrated into upper leaves then kept in standard light or dark (with infiltrated leaves covered in aluminum foil) for 24 hours. (B) GFP movement assays performed in *GUN5*- (TRV-*GUN5*) silenced plants as described for (A). (C) GFP movement assays performed in various plants with or without treatment with flg22 as described for (A). Statistical significance was measured by the Mann-Whitney U-test. * p<0.05, **p<0.005, ***p<0.0005, ****p<0.0001 (asterisks). ns = p>0.05. Error bars indicate +/- standard deviation.

Light can also determine the extent of PD-mediated intercellular trafficking [56, 57]. Indeed, this was a trend observed across all treatments (**Fig 5A** and **B**). PD-mediated trafficking was typically higher in dark-treated leaves compared to light-exposed leaves across treatments of non-silenced control plants (TRV-GUS), although the differences were not always statistically significant (**Fig 5A** and **5B**). However, a reverse trend was observed in *GUN5-*silenced plants, where PD-mediated trafficking was higher in the light than the dark across treatments.

### Flg22 depends on heme level to regulate PD-mediated trafficking

The biological role of ONPS and its potential CRSs like heme have not been clearly demonstrated. We therefore assessed the importance of heme-mediated ONPS in a plant defense response to a pathogen elicitor. For this we turned to the model bacterial elicitor, flagellin, and its flg22 peptide that is a potent inducer of pattern triggered immunity (PTI) [58]. Flg22-induced PTI results in decreased PD-mediated intercellular trafficking (Cheval et al., 2020). Flg22 has also been reported to induce the expression of *FC1* but reduce the expression of *FC2* [42]. To determine whether flg22 regulates PD-mediated trafficking in a heme-dependent manner, we infiltrated 500 nM flg22 into *GUN5*-silenced *N. benthamiana* plants, which had reduced heme levels. We found that treating a non-silenced control plant (TRV-*GUS*) with 500 nM flg22 significantly decreased intercellular trafficking (**Fig 5C**). The flg22 treatment did not decrease intercellular trafficking in *GUN5*-silenced plants compared to the mock treatment (**Fig 5C**). This lack of response to flg22 in *GUN5*-silenced plants suggests that there was insufficient heme to regulate intercellular trafficking in the silence plants. These findings point to a role for heme in facilitating the flg22-mediated regulation of intercellular trafficking, suggesting that this signaling pathway may have a wider role in plant-pathogen interactions.

### Identification of potential PD-associate nuclear genes (PDANGs)

We next set out to identify genes that regulate PD-mediated trafficking in response to defective chloroplast biogenesis and signaling mediated by TBS pathway intermediates. To understand which genes are involved in regulating PD-mediated trafficking by *ISE2* and *GUN5* and whether *ISE2* and *GUN5* utilize the same pathway for this regulation, we conducted a transcriptomic analysis by next-generation sequencing. In *ISE2*-silenced plants, 684 genes were significantly differentially expressed, with a cutoff for adjusted p-values of <0.05 and |log2FC| >0. This analysis revealed 269 genes whose expression was induced and 415 whose expression was repressed (**Fig 6A**). In the *GUN4-*silenced plants, only 18 genes were significantly differentially expressed, with nine genes induced and 9 suppressed. In striking contrast, in *GUN5-*silenced plants 1,573 genes were significantly differentially expressed, where expression of 809 genes was induced, and that of 764 genes were suppressed (**Fig 6A**). Only 8 differentially expressed genes (DEGs) were shared among *ISE2*, *GUN4*, and *GUN5*-silenced plants. A total of 241 genes were shared only between *ISE2* and *GUN5*-silenced plants, indicating a potential common regulatory mechanism (**Fig 6B)**. In terms of unique DEGs, *ISE2*-silencing resulted in 432 uniquely expressed genes, *GUN4* silencing presented only 1 unique gene, and *GUN5* silencing showed 1,318 unique genes, suggesting that silencing each gene has distinct gene-specific effects. The substantial overlap of shared genes between *ISE2* and *GUN5* suggests a potential functional link where ISE2-mediated RNA processing intersects with GUN5’s roles in plastid signaling. This implies that post-transcriptional regulation in plastids and tetrapyrrole biosynthesis are interconnected.

**Fig 6.**
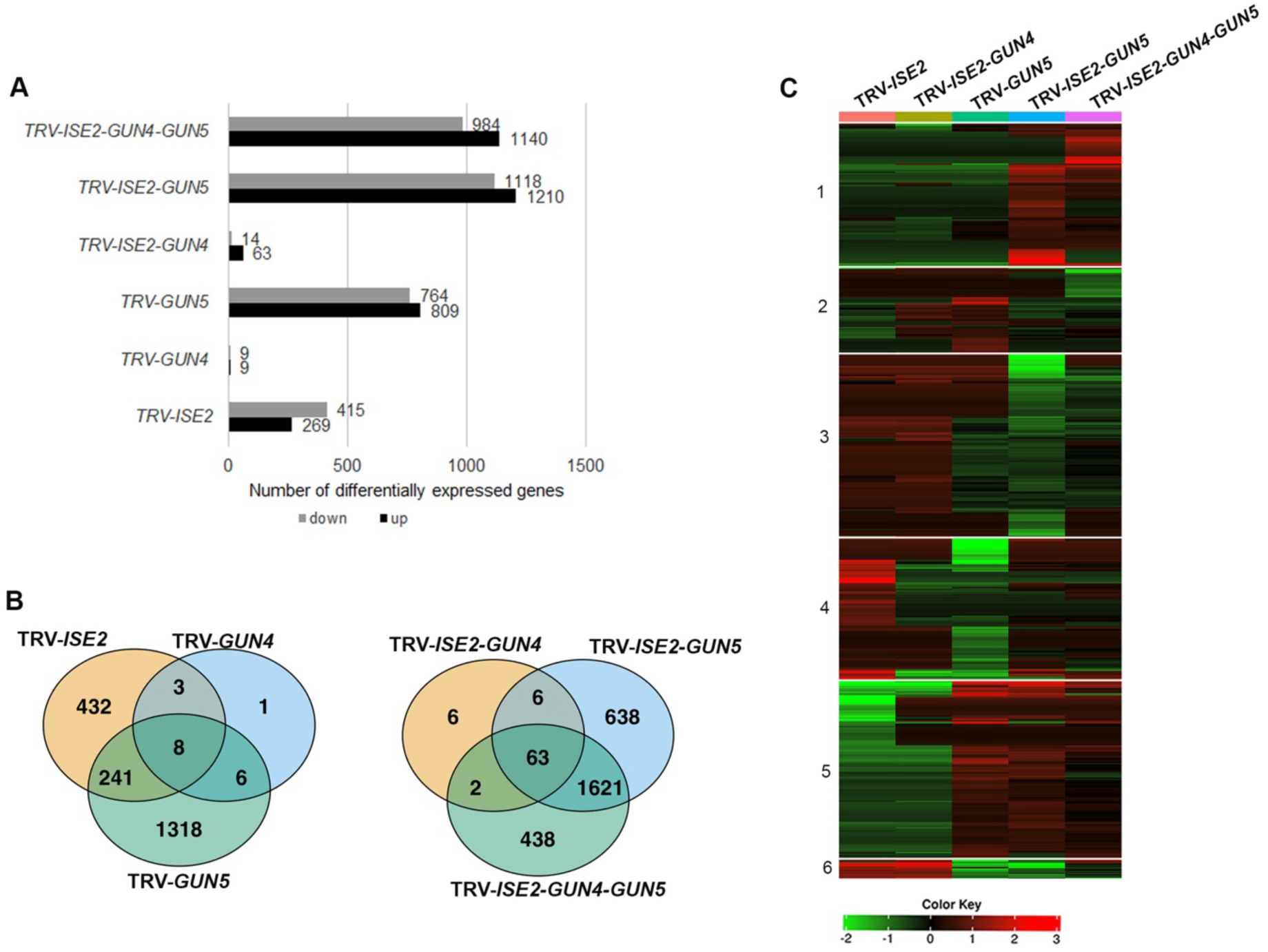
RNA sequencing analyses of *ISE2*-*GUNs*- silenced *N. benthamiana* plants. (A) Number of differentially expressed genes (DEGs) with adjusted p values below 0.05 in various silenced plants. All groups were compared to expression in non-silenced control plants (TRV-*GUS*). (B) Venn diagram showing the overlap of the DEGs in *ISE2-, GUN4-,* and *GUN5-* alone and *ISE2-GUN4-, ISE2-GUN5-, ISE2-GUN4-GUN5-* co-silenced plants. (C) DEGs of *ISE2-GUN-*silenced plants are clustered into six groups using k-means. Only DEGs with adjusted p values below 0.05 are shown, and values on the heat map show log_2_ fold change in expression.

For co-silenced plants, in *ISE2-GUN4* plants only 88 genes (63 induced and 14 suppressed) were differentially expressed. In contrast, in *ISE2-GUN5* and *ISE2-GUN4-GUN5* silenced plants 1,573 genes (809 induced and 764 suppressed) and 2,328 genes (1,210 induced and 1,118 suppressed) were differentially expressed, respectively. *ISE2-GUN4-co-*silenced plants showed 8 uniquely expressed genes, while *ISE2-GUN5-* and *ISE2-GUN4-GUN5-*silenced plants showed 638 and 438 expressed genes, respectively **(Fig 6B**). For shared DEGs, *ISE2-GUN4-* and *ISE2-GUN5-* silenced plants shared only 6 genes, whereas *ISE2-GUN5-* and *ISE2-GUN4-GUN5-* silenced plants shared 1,621 DEGs. This suggests that *GUN5-*silencing has a higher impact on differential gene expression in these co-silenced plants compared to silencing *ISE2- or GUN4*.

Clustering analysis of DEGs in heatmap following silencing of *ISE2-, GUN5-* alone and co-silencing of *ISE2-GUN4-, ISE2-GUN5-, ISE2-GUN4-GUN5-* revealed six clusters of patterns of gene expression **(Fig 6C)**. (DEGs from the *GUN4 alone*-silenced plants were omitted from further analyses because of the small number of genes). A striking observation was the opposing regulation of these clusters following the silencing of different genes alone or in combination. When *ISE2*, and *ISE2*-*GUN4* were silenced, DEGs in cluster 5 were repressed while the same DEGs were mostly induced after silencing *GUN5*, *ISE2-GUN5*, and *ISE2*-*GUN4*-*GUN5* **(Fig 6C)**. Conversely, DEGs in cluster 6 were upregulated in *ISE2* or *ISE2-GUN4* plants, while their expression was downregulated in *GUN5*, *ISE2-GUN5*, and *ISE2*-*GUN4*-*GUN5* silenced plants **(Fig 6C).** Gene ontology (GO) enrichment analysis revealed that DEGs in cluster 5 belonged to several cellular compartments, including cytoplasm, secretory vesicle, chloroplast, plasmodesma, and vacuole **(S1 Table)**. This suggests a link between chloroplast function and intercellular trafficking via PD. That DEGs in cluster 5 were involved in processes in several cellular compartments was supported by a KEGG analysis revealing broad effects on metabolism (**S2 Table**). In keeping with fewer DEGS belonging to cluster 6, only the plasma membrane was identified as a cellular compartment that was significantly affected in the GO analysis (**S3 Table**). Nonetheless, KEGG pathways that were significantly represented by the DEGs included the TBS pathway and other pathways including vitamin B6 and flavonoid metabolism, and terpenoid biosynthesis (**S4 Table**).

### Gene clusters associated with altered intercellular trafficking and heme metabolism

Focusing on clusters 5 and 6 (**Fig 6C**), we attempted to identify candidate PDANGs that were regulated by heme. After selecting genes belonging to the GO group plasmodesma-related terms, an in-silico pipeline, Plasmodesmata In silico Proteome 1 (PIP1) [59]. This pipeline compares four published PD proteomes which significantly overlap in terms of protein family and subfamily composition. The analysis output classifies whether the subfamily appears in one or multiple PD proteomes and is based on predicted distinctive features associated with PD-verified genes (235). The pipeline generates four lists of genes/proteins. Candidate list A includes genes encoding proteins from subfamilies found in more than one PD proteome with predicted signal peptide (SP), glycophosphatidylinositol anchor (GPI), or transmembrane domain (TM) features, which are the features enriched in PD-verified proteins. Candidate list B, while also appearing in multiple proteomes, does not have membrane-localizing features. Candidate lists C and D comprise proteins from subfamilies identified in a single proteome, with candidate list C including proteins with predicted targeting features and candidate list D including those without. Since experimentally verified PD proteins typically exhibit membrane-targeting features, candidate lists A and C are the most promising candidates. This analysis identified several genes encoded membrane-localized transporters and others from clusters 5 and 6 as potential PD genes regulated by heme (**Table 1**). A tetraspanin was also among these potential heme-regulated genes, although this prediction was made with less confidence than the others.

**Table 1.** Predicted candidate PD genes from PD-enriched GO terms in clusters 5 and 6 using the Plasmodesmata In silico Proteome 1 (PIP1) pipeline.

| *Nicotiana benthamiana*<br>gene ID | TAIR ID<br>(Arabidopsis<br>ortholog) | Protein names | Candidate list<br>or known PD<br>gene |
| --- | --- | --- | --- |
| *Niben261Chr07G1355001* | *AT1G11260* | Sugar transport protein 1 (Glucose transporter) (Hexose transporter 1) | A |
| *Niben261Chr17G0890004* | *AT5G26340* | Sugar transport protein 13 (Hexose transporter 13) (Multicopy suppressor of snf4 deficiency protein 1) | A |
| *Niben261Chr08G0042012* | *AT3G13750* | Beta-galactosidase 1 (Lactase 1) (EC 3.2.1.23) | A |
| *Niben261Chr08G1069003* | *AT4G27500* | Proton pump-interactor 1 | A |
| *Niben261Chr05G0827010* | *AT2G38290* | Ammonium transporter 2 (AtAMT2) | A |
| *Niben261Scf01608G0000028* | *AT4G11820* | Hydroxymethylglutaryl-CoA synthase (HMG-CoA synthase) (EC 2.3.3.10) (3-hydroxy-3-methylglutaryl coenzyme A synthase) (Protein EMBRYO DEFECTIVE 2778) (Protein FLAKY POLLEN 1) | B |
| *Niben261Chr11G1196005* | *AT1G45145* | Thioredoxin H5 (AtTrxh5) (Protein LOCUS OF INSENSITIVITY TO VICTORIN 1) (Thioredoxin 5) (AtTRX5) | B |
| *Niben261Chr03G1819003* | *AT1G09640* | Probable elongation factor 1-gamma 1 (EF-1-gamma 1) (eEF-1B gamma 1) | B |
| *Niben261Chr16G0270015* | *AT4G27450* | AT4g27450/F27G19_50 (Aluminum induced protein with YGL and LRDR motifs) | B |
| *Niben261Chr03G0692005* | *AT4G16130* | L-arabinokinase (AtISA1) (EC 2.7.1.46) | B |
| *Niben261Chr15G0702022* | *AT5G22060* | Chaperone protein DnaJ 2 (AtDjA2) | B |
| *Niben261Chr06G1362001* | *AT5G20250* | Probable galactinol--sucrose galactosyltransferase 6 (EC 2.4.1.82) (Protein DARK INDUCIBLE 10) (Raffinose synthase 6) | B |
| *Niben261Chr13G0053007* | *AT4G34150* | At4g34150 (Calcium-dependent lipid-binding (CaLB domain) family protein) | B |
| *Niben261Chr13G1069004* | *AT5G62740* | Hypersensitive-induced response protein 1 (AtHIR1) | B |
| *Niben261Chr07G0225014* | *AT5G44800* | Protein CHROMATIN REMODELING 4 (AtCHR4) (EC 3.6.4.-) (Protein PICKLE RELATED 1) | B |
| *Niben261Chr10G1117004* | *AT5G60220* | Tetraspanin-4 | C |
| *Niben261Chr03G1126002* | *AT2G28790* | Pathogenesis-related thaumatin superfamily protein | Not |

To extend the identification of PDANGs potentially regulated by heme and the TBS pathway, we manually extracted DEGs encoding cell wall biosynthesis and modification enzymes as well as DEGs encoding known PD-associated proteins (**Fig 7**). The expression of cell-wall related genes was almost entirely unaffected by silencing *ISE2* alone. In contrast, expression of numerous genes related to cell-wall synthesis and modification was altered in leaves of plants where GUN5 was silenced alone or together with *ISE2* or *ISE2* and *GUN4* (**Fig 7**). Notably, expression of callose and cellulose metabolism genes was mostly repressed in *GUN5*-silenced plants with reduced heme while expression of xyloglucan-related genes were upregulated in the same samples. The suggested involvement of the xyloglucan modifying XYLOGLUCAN ENDOTRANSGLYCOSYLASE/HYDROSYLASE (XTH) enzymes supports previous findings linking this class of cell wall carbohydrates to altered PD-mediated intercellular trafficking [12].

**Fig 7.**
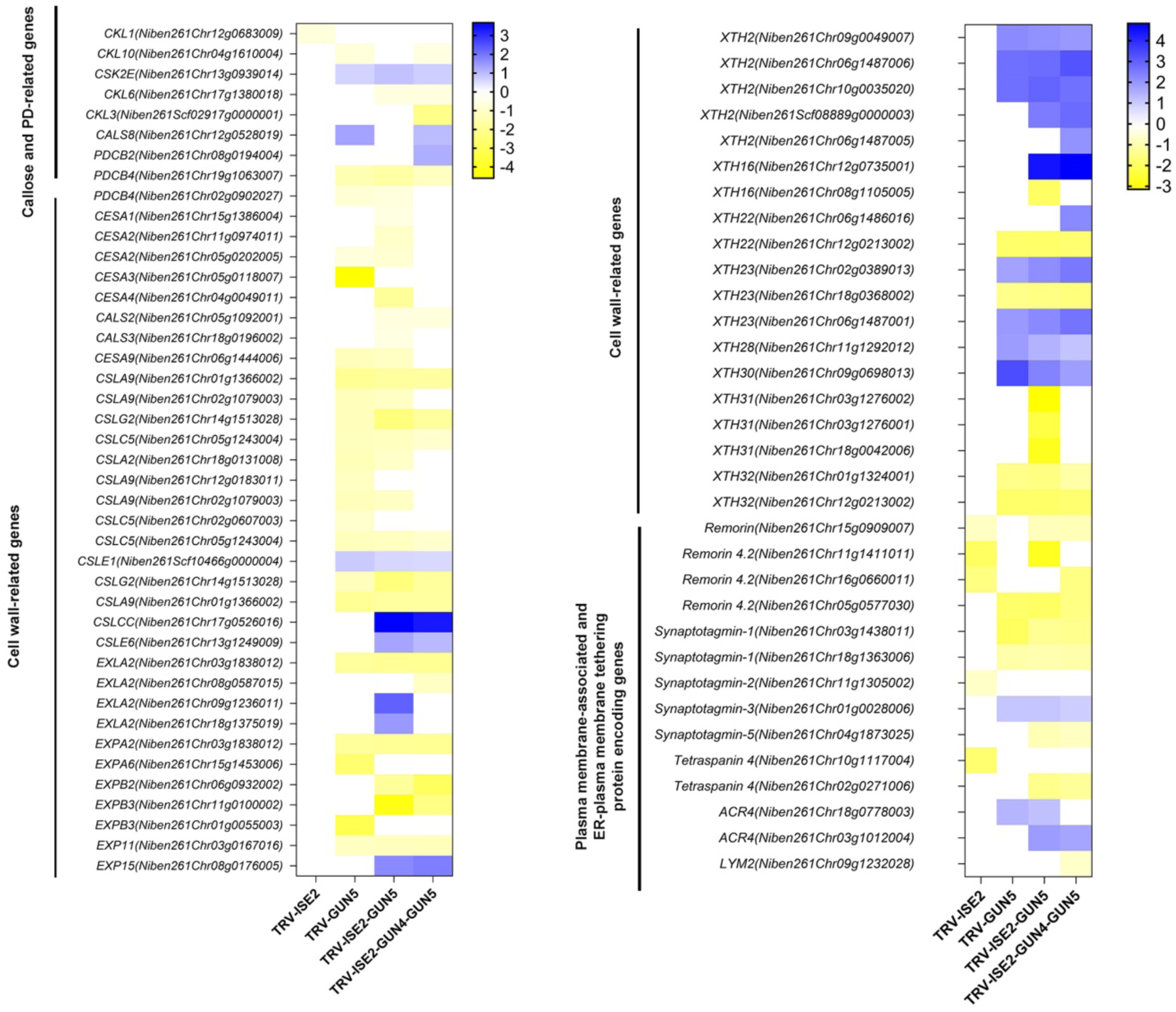
Heatmap showing expression of callose-, PD-, cell wall-related DEGs in *N. benthamiana* plants where ISE2 and/or GUN5 was silenced. DEGs related to callose metabolism, cell wall synthesis and modification, and PD structure and function were identified by RNA-seq analysis. Only DEGs with adjusted p values below 0.05 are shown, and values on the heat map show log_2_ fold change in expression.

While expression of most of the cell wall-associated genes was unaffected by silencing *ISE2*, expression of several *Remorin*, a *SYNAPTOTAGMIN* and a *TETRASPANIN* genes was repressed. Remorins are plant-specific proteins that localize to PD and to the PM where they are markers for membrane microdomains (lipid rafts) [60]. Remorins are important for determining the outcome of specific plant-virus interactions, supporting their roles in PD functions (reviewed in [61]). *Remorins* were also differentially expressed in plants where GUN5 alone, *ISE2-GUN5* and *ISE2* along with *GUN4* and *GUN5* were silenced, although the specific gene silenced differed for each group of plants (**Fig 7**). SYNAPTOTAGMINs (SYTs) regulate calcium-dependent membrane fusion, endo- and exocytosis, and have roles as tethers for ER-PM contact sites [62]. While expression of several *SYTs* responded to silencing GUN5 in various contexts, one *SYT3* gene (*Niben261Chr01g0028006*) apparently responded to any silencing of GUN5 but not ISE2 (**Fig 7**), making it an interesting candidate for follow up studies. Tetraspanins are scaffolding proteins which are wildly distributed in plants and animals [63, 64]. They have been identified as core plasmodesmal proteins and have been proposed to serve essential roles in PD [65]. Notably, expression of *TETRASPANIN 4* homeologs was repressed in all instances where *ISE2* was silenced (**Fig 7**), suggesting that it might be a key component of the response to signaling initiated when chloroplasts lack ISE2. ARABIDOPSIS CRINKLY4 (ACR4) is a PD-localized receptor kinase with a role in maintaining the root meristem, likely through complex formation with CLAVATA1 [66]. Expression of *ACR4* genes was induced when *GUN5* was silenced, making them possible mediators of the *GUN5* silencing phenotype of reduced intercellular trafficking.

### Plasmodesmal density not callose level correlates with heme levels and intercellular trafficking

Defects in intercellular trafficking can be associated with changes in PD density [20, 46]. Arabidopsis PLASMODESMATA-LOCATED PROTEIN1, a general marker for PD [67], tagged with GFP (PDLP1-GFP) was expressed in the leaves of silenced to measure PD distribution. PD are nanopores with an average diameter of less than 30 nm, so we assume that the GFP foci in the cell wall observed by confocal microscopy are likely clusters of PD since individual PD are beyond the resolution limits of confocal microscopy. The average number of clusters (per 100 µm^2^ cell wall) was higher in leaves of *ISE2*-silenced plants, as expected based on previous findings, although this number was not statistically significant in this group of experiments. Consistent with the results of earlier analysis, PD density was also increased in *ISE2-GUN4* co-silencing plants which also had increased intercellular trafficking (**Fig 2B**). No change in PD density was measured in leaves where only *GUN4* was silenced (**Fig 8A**), in agreement with no changes with intercellular trafficking (**Fig 2B**). In all plants where *GUN5* was silenced, whether individually or with other genes, PD density was significantly lower than in non-silenced control (TRV-*GUS*) plants (**Fig 8A**). These results support a positive correlation between plasmodesmal density and the extent of intercellular trafficking.

**Fig 8.**
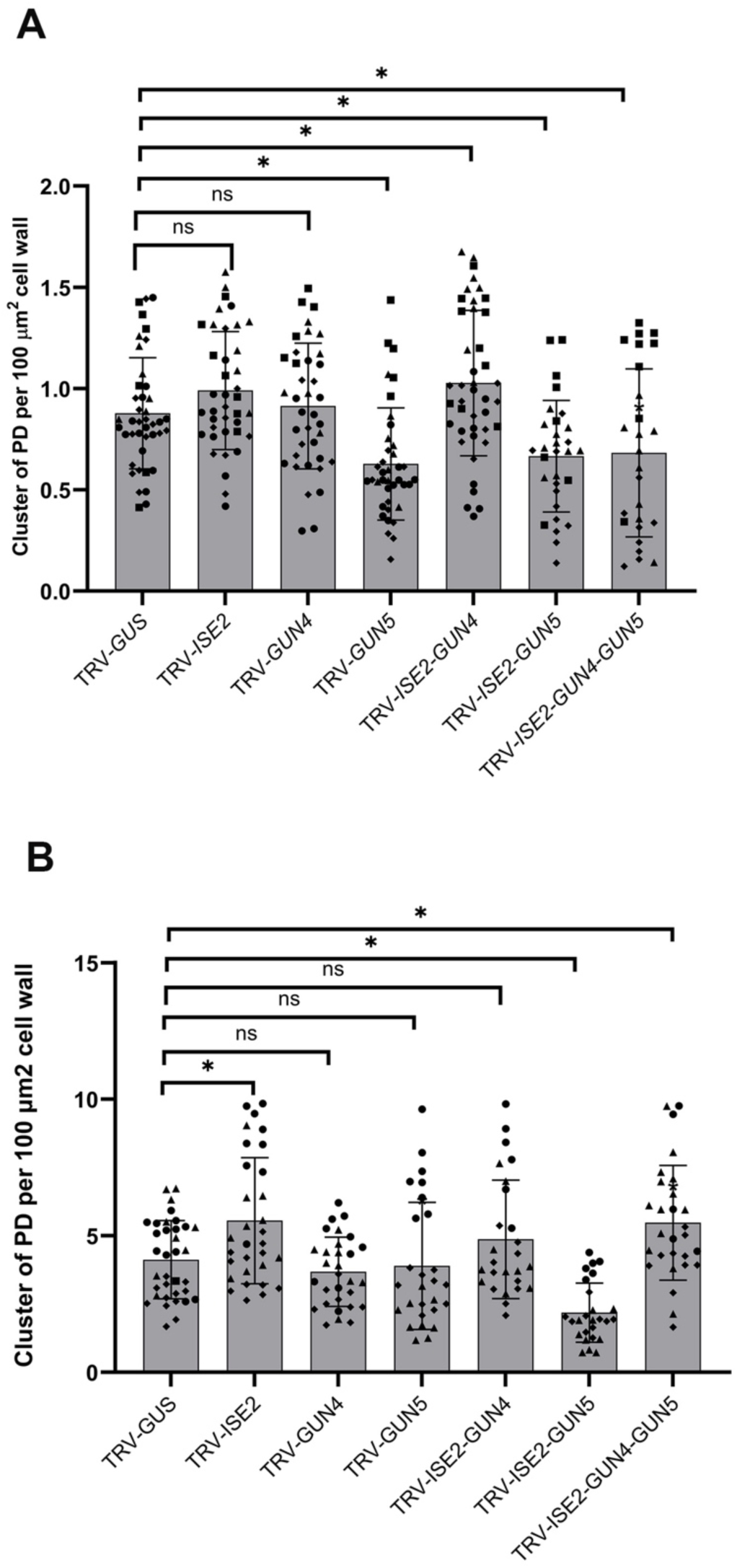
PD density is correlated to intercellular trafficking in *N. benthamiana* in response to heme. (A) Clusters of PD were labeled with Arabidopsis PD marker PDLP1-GFP. Images were collected from z-stacks and density per unit cell wall area (100 µm^2^) was calculated. (B) Aniline blue staining was used to stain callose and detect clusters of PD. Images were collected from z-stacks and density per unit cell wall area (100 µm^2^) was calculated. For (A) and (B), outliers identified interquartile range (IQR) analysis were excluded. Student’s t-test was used to determine statistical significance compared to non-silenced controls (TRV-*GUS*), asterisks indicate p<0.05. ns= p>0.05. Error bars indicate +/- standard deviation.

Callose accumulation in the cell wall surrounding PD can impair intercellular trafficking, and this effect is alleviated when callose is removed [10]. To gauge the relationship between callose and the intercellular trafficking observed in silenced plants, leaves from silenced plants were stained with aniline and the number of stained foci scored as described [68]. While the number of aniline-blue foci was increased in leaves of *ISE2*-silenced plants compared to those of non-silenced control plants, there was no change in *ISE2-GUN4* co-silenced plants (**Fig 6B**). This observation was incongruent with the increased intercellular trafficking measured in both groups of silenced plants (**Fig 2B**). Similarly, there was no change in the number of aniline-blue stained foci in the leaves of *GUN5*-silenced plants but decreased number of foci *ISE2-GUN5* co-silenced plants and increased numbers of foci in leaves of *ISE2-GUN4-GUN5* co-silenced plants (**Fig 6B**). However, intercellular trafficking was decreased in all plants where GUN5 was silenced or co-silenced (**Fig 2B**). These results do not indicate a correlation between the number of aniline-blue stained foci and the propensity for intercellular trafficking. Considering the findings of these two assays, it is likely that changes in PD density may be responsible for the changes in intercellular trafficking observed.

## DISCUSSION

Heme is an essential biomolecule that transfers electrons and catalyzes numerous biochemical reactions. It has been proposed that heme was likely used by the earliest forms of cellular life in the ferrous iron (Fe^2+)^-rich environments in which primordial life evolved [69]. The utility of heme has had rendered it essential for energy, redox metabolisms, and the synthesis of coenzymes and pigments. It acts as a prosthetic group for many proteins involved in essential biological processes such as respiration, photosynthesis, and the metabolism and transport of oxygen [70]. Heme is synthesized in most organisms through a highly conserved biosynthetic pathway [69]. Heme binds Fe^2+^ through four nitrogen atoms in its protoporphyrin ring, functioning as a prosthetic group in various apoproteins. The main type, heme b, is synthesized from protoporphyrin IX by ferrochelatase (FC) [71, 72]. Other heme forms, including a and c, derive from heme b. Heme a and b attach non-covalently to hemoproteins, while heme c is covalently linked to cysteine residues, as in cytochrome C. Beyond a prosthetic group, heme also serves as a regulatory molecule, with free heme produced by FCs for metabolic processes [45]. Additionally, enzymes like catalase, peroxidase, or cytochrome P450 rely on heme as a critical cofactor. Additionally, heme plays roles in influencing transcription [73–77], translation [78], post-translational modification [79], and ion-channel function [80].

In this study, we showed that heme can modulate PD functions, reinforcing the view that heme is not confined to organelle-localized activity but acts as a multi-functional signal that integrates chloroplast and nuclear processes to modulate PD functions. Our results reveal a link between two ancient and foundational features of plants. Given that PD are crucial to the evolution of multicellularity and are essential for plant survival [43], this regulation by heme may have been crucial in the evolution of multicellular plant systems when the coordination of growth, development, and response to the environment became essential. Interestingly, the SAL1–PAP CRS pathway is involved in regulating stomata in streptophyte algae (a sister clade to modern land plants), suggesting their relationship is ancient [81]. We therefore propose a similar ancient relationship between heme and PD. This framework redefines chloroplasts—originating from endosymbiotic cyanobacteria—as central hubs for both energy metabolism and signaling, offering a mechanistic basis for understanding how retrograde and symplastic pathways integrate to regulate plant development and responses to environmental changes. In plants and algae, heme is exclusively made in chloroplasts, so it may be a general indicator of chloroplast health and number and serves as a proxy for photosynthetic and other metabolic output. By using heme to regulate intercellular trafficking, the plant can coordinate the production and distribution of metabolites between cells and tissues via plasmodesmata.

### The role of TBS pathway and heme in regulating intercellular trafficking

The *<u>g</u>enome <u>un</u>coupled* (*gun*) mutants have defective CRS, resulting in the deregulation of expression of PHANGs [26, 29]. Here we show that at least one of these mutants, *gun5*, also has deregulated intercellular trafficking via PD (**Fig. 1B**). Silencing the *N. benthamiana GUN*5 homoeolog also perturbed intercellular trafficking (**Fig. 1D** and **1E**). While silencing or knockout of *GUN5* disrupted intercellular trafficking, similar changes in *GUN4* expression had no effect on PD-mediated intercellular trafficking (**Fig. 1D** and **1E**), pointing to the enzymatic activity of the Mg-chelatase as a critical control point for generating signals that regulate PD permeability. Chloroplast RNA metabolism gene *ISE2* was found to regulate PD-mediated trafficking in *GUN5* dependent pathway. Consistent with this, reduced heme levels were present in other plants in which decreased intercellular trafficking was reported (TRV-*RH3, RNaseJ, PNPAseA/B, IPI1;* [20]; **Fig 2F**). An interesting and unexpected observation of this study was that exogenous heme and ALA can increase PD-mediated trafficking (**Fig 4**), and heme exerted its effects independent of light (**Fig 5**). It is not clear that application of heme increases intracellular heme concentrations, but in the context of our experiments an effect was apparent. The use of novel tools like fluorescent reporters for the subcellular measurement of free heme could be useful in future studies to probe the response to exogenous heme application [82]. The connection between PD and Mg-chelatase is supported by the observed correlation between the density of clusters of PD and PD-mediated trafficking, where decreased density of clusters of PD was measured in plants with decreased intercellular trafficking and vice versa (**Fig. 1F and 2D**).

### Threshold heme levels may regulate intercellular trafficking

Heme is reported to translocate from the chloroplast to the nucleus, where it regulates transcription, RNA metabolism, and chromatin remodeling [48]. Heme-related signals are proposed to be the primary mediator of CRS during chloroplast biogenesis [83]. Heme’s low solubility in water suggests it predominantly exists either dissolved in membranes or non-covalently attached to proteins [84]. Thus, free heme is defined as heme that was synthesized by the FCs which is available for further metabolic conversion [85]. When the chloroplast is disrupted by norflurazon, an inhibitor of carotenoid biosynthesis, total heme levels are reduced, but endogenous free heme levels are increased [85]. However, free bulk heme levels do not neatly correlate with the *gun* phenotype, suggesting that if heme acts as an organelle-derived signal, its transfer likely occurs through specific transport mechanisms and protein interactions [47]. Indeed, our results suggest that bound heme levels are important for regulating intercellular trafficking.

In *N. benthamiana,* we found that *GUN5*-silenced plants showed decreased total heme content. Papenbrock et al. [49] also reported that transgenic tobacco (*N. tabacum*) with reduced levels of *CHLH* expression had reduced heme levels, which they attributed to negative regulation of ALA synthesis. Nonetheless, decreased total heme content may affect CRS, which may in turn regulate the expression of PD-related and/or cell-wall remodeling genes to modulate intercellular trafficking via PD (**Fig 7** and **8)**. Silencing *ISE2, GUN4* alone, and the combination *ISE2-GUN4* does not significantly impact heme levels compared to the non-silenced control. These silenced plants either increase or do not affect PD-mediated trafficking **(Fig 1** and **2)**. In contrast, silencing *GUN5, ISE2-GUN5*, and *ISE2-GUN4-GUN5* caused reduced levels of heme and decreased PD-mediated trafficking. Thus, reduced heme levels correlate with decreased intercellular trafficking, indicating that heme is essential for normal trafficking. A nuanced interpretation of our findings is that intercellular trafficking may depend on maintaining a certain heme threshold level. If heme levels fall below this threshold (e.g., in *GUN5 or ISE2-GUN5* co-silenced plants), trafficking is decreased. At or above this threshold, trafficking remains stable or can be increased depending on other factors or signals. The reduced heme levels in *GUN5* and other silenced plants may influence the demand for heme in cytochrome b_6_f, affecting electron transfer in the photosynthetic machinery and may generate reactive oxygen species (ROS). Therefore, the role of redox signaling and other CRS in the regulation of trafficking via PD cannot be ruled out. Indeed, our results suggest that heme alone may not be sufficient to increase intercellular trafficking as they do not fully explain the increased trafficking observed in *ISE2-, RH39-* or *PNPaseA*-silenced plants [20].

### Distinct role of two different pools of heme in regulating PD-mediated trafficking

Here, we show that reducing FC2, but not FC1, activity leads to reduced total heme and intercellular trafficking, suggesting that FC2 may be producing a specific pool of heme involved in PD function (**Fig 3B** and **3C**). FC2 appears to produce most of the heme used by the photosynthetic machinery whereas FC1 may mainly produces heme that is exported outside the chloroplast for housekeeping functions [42]. However, more recent work has suggested that such a simple separation of functions is unlikely. For instance, promoter swap experiments have demonstrated that FC2 can mostly compensate for a complete lack of FC1 protein, suggesting that at least some heme produced by FC2 can be exported from chloroplasts [86]. Our results support this hypothesis and further suggest that this FC2-derived pool of heme has signaling functions for regulating PD and PDANG expression. Interestingly, this pool appears distinct from the one produced by FC1 that can signal to control PhANG expression [27]. Thus, the signaling roles of heme and specific heme pools is more complicated than previously thought and will require further investigation to understand their separate mechanisms.

Disrupting heme production in chloroplasts can also affect the supply to other organelles, such as mitochondria and peroxisomes [87]. In mitochondria, heme is essential for cytochrome *c* in the mitochondrial electron transport chain to transfer electrons critical for ATP production. In peroxisomes, heme acts as a co-factor for peroxidase and catalase in redox signaling. We cannot rule out the possibility of heme acting through other organelles to regulate the PD-mediated trafficking. For instance, heme might be transported to mitochondria/other cellular organelles or directly to the nucleus to regulate the expression of PD/cell-wall-associated nuclear genes to regulate PD-mediated trafficking. Notably, the mitochondria-localized protein, ISE1, can regulate intercellular trafficking via PD [88]. Since chloroplast-produced heme is required for mitochondrial electron transport (mETC), reduced total heme level may affect mitochondrial function, which could regulate the trafficking through PD.

### Heme regulation of plasmodesmata is not light dependent

In control plants, intercellular trafficking was found to depend on light conditions (**Fig. 5**), as has been reported previously [56, 57]. In the dark, reduced ATP levels due to reduced photosynthesis could open the PD for the export of the assimilated carbon from the leaves [89]. *GUN5*-silenced plants did not show increased trafficking in dark conditions, indicating that endogenous heme or tetrapyrrole signaling is necessary for this regulation. Exogenous heme modified PD-mediated trafficking in plants with an intact TBS pathway independently of light, supporting a direct role for heme in signaling and ruling out the likelihood of signaling caused by photo-oxidative damage (**Fig 5A).** In contrast, heme treatment of *GUN5*-silenced plants increased trafficking only in light (**Fig 5B**). Thus, the introduced heme can modulate PD function, possibly through redox signaling, callose regulation, or interactions with PD-associated proteins. Additionally, GUN5 may play a crucial role in integrating light cues into PD function, potentially through a retrograde signaling pathway.

## MATERIAL AND METHODS

### Plant Materials and Growth Conditions

*N. benthamiana* seeds were sown in pots filled with hydrated soil (Berger 35). After 10 days, germinated seedlings were transplanted into individual pots and allowed to grow for 10-12 days in a 16-hour photoperiod before performing VIGS. Table **S5** lists the *Arabidopsis thaliana* ecotype Columbia (Col-0) and *gun* mutant lines used in this study. These mutant lines are in the Col-0 background and have previously been described [26]. Arabidopsis seeds were also sown directly in pots and stratified for 3 days at 4°C before moving to a growth chamber with a 16-8-hour photoperiod of 120 μmol m^-2^ · s^−1^ at 23°C until further experimentation. For all experiments samples were collected 3-5 hours after the onset of the light period.

### RNA isolation, cDNA synthesis and quantitative polymerase chain reaction (qPCR)

The ninth/tenth leaf (counted from the bottom) of silenced or non-silenced *N. benthamiana* plants was collected and flash-frozen in liquid nitrogen. RNA was isolated from three independent samples using the RNA isolation kit (Qiagen), following the manufacturer’s instructions. To remove DNA, the RNA was treated with DNase using the DNA-free DNA removal kit (Invitrogen, CA) for subsequent qPCR or RNA sequencing analysis. Complementary DNA (cDNA) was synthesized with oligo-dT primers and M-MLV reverse transcriptase (Promega, WI), following the manufacturer’s instructions. cDNA was then used for gene cloning or quantitative polymerase chain reaction (qPCR). Diluted cDNA (1:5) with nuclease-free water is used for qPCR to quantify the expression of photosynthesis-associated nuclear genes (PHANGs) and to confirm the silencing in *VIGS*-infiltrated plants. qPCR was performed using the LightCycler 480 SYBR Green I Master Mix and a LightCycler 480 Real-Time PCR system (Roche, Indianapolis, IN, USA) with the following thermal cycling program: 95°C for 10 minutes, followed by 40 cycles of 95°C for 10 seconds, 59°C for 10 seconds, and 72°C for 20 seconds. The expressions of PHANGs and silenced genes are normalized to the expression of *EF1-α*.

### Gene cloning to generate Virus-Induced Gene Silencing (VIGS) constructs

The *Arabidopsis thaliana genome uncoupled* (*GUN*) loci were obtained from The Arabidopsis Information Resource database (TAIR, https://www.arabidopsis.org/index.jsp). These gene loci were *AtGUN2* (*AT2G26670*), *AtGUN4* (*AT3G59400*), and *AtGUN5* (*AT5G13630*). The protein sequences of each locus were used to obtain the corresponding *N. benthamiana* gene sequences (https://nbenthamiana.jp/). The gene loci in *N. benthamiana* are *NbGUN4* (*Nbe.v1.1.chr09g39610, Nbe.v1.1.chr16g37380*), *NbGUN5 (Nbe.v1.1.chr04g34700, Nbe.v1.1.chr14g15590*), *NbFC1* (*Nbe.v1.1.chr10g28700, Nbe.v1.1.chr17g40820*), *NbFC2* (*Nbe.v1.1.chr09g34020, Nbe.v1.1.chr11g29350*). The *NbGUN2* homoeologs were previously identified [90]. Gene fragments, approximately 400 bp in length, were amplified from cDNA from wild-type *N. benthamiana* plants using Phusion DNA polymerase (New England Biolabs, Ipswich, MA), and primers and plasmids are listed in **Table S6** and **S7,** respectively. The amplified silencing fragments were cloned into the pTRV2 (pYL156) vector (containing TRV RNA2, [91]) after digestion with restriction enzymes (New England Biolabs). To generate a co-silencing construct with *ISE2*, the *GUN4* and/or *GUN5* silencing fragments were digested and ligated to pTBS136, which already carried a fragment for silencing *ISE2*. Ligations were carried out with T4 DNA ligase (New England Biolabs). The resulting ligated plasmids were transformed into chemically competent DH5α *Escherichia coli* cells and cultured on LB media containing kanamycin for selection. Positive clones were screened through colony PCR. Plasmids were isolated from overnight-cultured positive clones using a mini-prep kit (QIAGEN), and the nucleotide sequences of silencing constructs in plasmid were confirmed through Sanger sequencing. These positive plasmids containing the silencing constructs were transformed into *Agrobacterium tumefaciens* GV3101 competent cells by chemical transformation (freeze/thaw) and used for VIGS experiments.

### VIGS

VIGS was performed as previously described [91]. Briefly, pYL192 (which expresses TRV1) and pTRV2 with the silencing construct were cultured overnight, centrifuged, and resuspended in an infiltration buffer containing 10 mM MgCl_2_ and 2-(N-morpholino) ethane sulfonic acid (pH 5.5, adjusted with KOH), along with 200 µM acetrosyringone, (Sigma, St Louis, MO, USA) to an optical density (OD) of 0.1 at 600 nm. An equal volume of pTRV1 and pTRV2 derivatives were mixed before infiltrating into the leaves of three-week-old plants. A TRV2 vector a fragment of GUS [12] was used as a negative control for all VIGS experiments. Movement assays were conducted 10-12 days after VIGS infiltrations.

### GFP-Movement Assay and VIGS

Intercellular trafficking in *N. benthamiana* was measured by a GFP movement assay, as described previously [20]. In brief, Agrobacteria carrying a binary construct for GFP expression (*p35S: GFP*) were cultured overnight in LB medium with antibiotics at 28°C. The cultures were centrifuged at 3000 rpm for 15 minutes, washed, and then resuspended to an OD of 0.0001 at 600 nm in infiltration buffer. To induce virulence, the Agrobacteria were incubated at room temperature for 3 hours with gentle shaking before infiltrating them into silenced leaves using a needleless syringe. Imaging was performed 48 hours post infiltration using a Leica SP8 confocal microscope (Leica Microsystems, Buffalo Groove, IL) with excitation and emission wavelengths of 488 nm and 500-530 nm, respectively. Images were collected using 20X objective with the pinhole aperture was set to 1 Airy unit. Z-stacks of GFP foci were maximum projected and analyzed using ImageJ [20]. The number of rings of cells in the epidermis surrounding the original GFP-expressing cell was determined. At least three biological replicates, each including at least 20 foci, were performed.

To measure intercellular trafficking in Arabidopsis *gun* mutant lines, the fifth and sixth leaves from five three-week-old plants were collected and low-pressure (60 psi) bombarded with plasmid pRTNL2rsGFP for the GFP movement assay as described [92]. After bombardment, the leaves were incubated abaxial side down on half-strength MS media at 23°C in a growth chamber. After 24 hours of bombardment, confocal microscopy was performed as described above to measure the GFP movement. At least 15 foci per sample were collected to analyze the GFP movement, and experiments were performed for at least three biological replicates. For statistical analyses, both the Mann-Whitney U test and the Bootstrap method [93] were performed to compare the GFP movement between samples.

### Quantification of PD density and staining callose with aniline blue

Clusters of PD in *N. benthamiana* plants were quantified after transiently expressing Arabidopsis PDLP1 fused to GFP (p35S: PDLP-GFP) as described previously [20]. Briefly, *Agrobacterium* containing PDLP-GFP was infiltrated at OD_600 nm_= 0.001. After 72 hours post infiltration, 15 um z-stacks of epidermal cell images at a step size of 1 um were collected with a Leica SP8 confocal microscope with a White Light Laser (Leica Microsystem, IL, USA) using a 25x water emersion objective. Three biological replicates, each consisting of sixty foci, were collected for each sample. Maximum intensity projections were generated for each of the z-stacks using ImageJ. Cell wall lengths were measured by tracing the cell wall in the field of view. Images were then converted to 8-bit images and inverted. After adjusting the intensity threshold, the particle analyze function was used to count the total number of punctate in the image. PD pitfields are measured per 100 µm^2^ cell wall area (cell length was multiplied by z-stacks size to measure the cell wall area). Mann-Whitney U-test was performed to measure the statistical significance comparing the PD pitfields between silenced and non-silenced control plants.

Callose at PD was stained using aniline blue and quantified using Method 1 described previously [68]. Images were obtained by exciting the aniline blue with a 405 nm laser, and emission was captured from 415 to 525 nm using a 40X water immersion objective. Z-stacks were acquired using a system-optimized step size of less than 1 μm. In total, 60 foci/regions of interest were collected for each sample across three biological replicates.

### Flg22 treatment, ALA feeding, and heme feeding

The stock solutions of flg22 (GenScript, Piscataway, NJ, USA, Cat. No. RP19986) and ALA (Sigma-Aldrich Inc., Cat. No. A3785) were freshly prepared in sterile water. Flg22 was further diluted to the desired concentration (0 nM, 100 nM, and 500 nM) using water. ALA was diluted to the desired concentrations (0 mM, 0.2 mM, 05 mM, 1 mM, 2 mM, 5 mM) using 10 mM phosphate buffer and 5 mM MgCl_2_. The stock solution of heme (100 mM) was prepared in DMSO and further diluted freshly to different concentrations in 100 mM KOH.

For the flg22 treatment, freshly prepared flg22 at desired concentrations were syringe-infiltrated into the leaves of *GUN5-* and non-silenced plants seven hours before confocal microscopy. For ALA and heme feeding, the desired concentrations of heme were infiltrated into *GUN5-* and non-silenced control plants 24 hours before confocal microscopy. Plants were kept in the dark overnight and moved to light for another 12 hours. For the GFP movement assay, these leaves were infiltrated with Agrobacterium carrying *35S:GFP* 48 hours before confocal microscopy. Movement assays were performed as described previously.

### Heme measurements

Non-covalently bound heme was measured in plants 10-12 days post VIGS infiltration. Ten leaf discs were collected from the eighth or ninth leaf of a plant and weighed to determine fresh weight. 100 mg of plant tissue was ground in liquid nitrogen using a mortar and pestle and the powder was dissolved in 1 mL of ice-cold extraction solution 1 (90% (v/v) acetone and 10% (v/v) 0.1 M NH_4_OH) and centrifuge at 10,000 x g for 10 minutes and the supernatant was discarded. The pigment extraction steps were repeated four times. To extract heme from the pellet, 500 µL of extraction solution 2 (80% (v/v) acetone, 16% (v/v) DMSO, and 4% (v/v) concentrated HCl) was added and mixed thoroughly. The solution was incubated for 10 minutes at room temperature and centrifuged at 10,000 x g for 10 minutes. The supernatants were diluted 100 to 1,000-fold with 10 mM KOH, and 10 µL of diluted samples were used for heme quantification.

Heme extracts were diluted in 10 mM KOH before being applied to the Horse radish peroxidase (HRP) enzyme assay. The HRP apoenzyme spontaneously reconstitutes with heme, resulting in an active enzyme [94]. The peroxidase activity of HRP (Biozyme) is utilized to oxidize luminol in the presence of H_2_O_2_, and the resulting luminescence signal is proportional to the amount of heme present in the sample. To prepare the master mix for each reaction, 50 µL of 2x reaction buffer, 30 µL of H2O, and HRP were added to achieve a final concentration of 2.5 nM. Eighty µL of the master mix was incubated with 20 µL of the sample or standards for 30 minutes at room temperature, followed by the addition of the chemiluminescent western blot detection reagent (Pierce^TM^ ECL, ThermoScientific, Rockford, IL), prepared according to the manufacturer’s instructions (50 µL of luminol and 50 µL of peroxide solution), and incubated for 12-15 minutes. Finally, the luminescence was measured at 430 nm using a 96-well plate reader in luminescence mode with an integration time of 0.5 seconds. Two standard curves of 250-4000 nM and 50-400 nM were used to quantify total and chloroplast heme. Chloroplasts were isolated as described [95].

### Immunoblots

Ninth/tenth leaves of the non-silenced or silenced plants were harvested and ground in liquid nitrogen with a mortar and pestle to a fine and homogeneous powder. 300 mg of this powder was transferred to a new microcentrifuge tube, and 500 μL of extraction buffer (containing 150 mM NaCl, 20 mM Tris-HCl pH 7.5, 1 mM EDTA, 1% Triton X-100, 0.1% β-mercaptoethanol (ME), protease inhibitor, and 1 mM phenylmethylsulfonyl fluoride (PMSF) was added. The samples were immediately mixed on a vortex and were centrifuged at maximum speed at 4 °C for 10 minutes. The supernatants were transferred to a new tube and centrifuged again for 5 minutes at full speed at 4 °C. A total protein extract (36 µL) was then loaded onto a 10% SDS-PAGE gel. The resolved proteins were transferred to a polyvinylidene difluoride (PVDF) membrane (Millipore, Billerica, MA, USA) and probed with the indicated antibodies at specified dilutions. The primary polyclonal antibodies (Agrisera) targeting the specified proteins were used at these dilutions: Cytb6 at 1:40,000, Cytf at 1:20,000, D1 at 1:20,000, and Tubulin at 1:1,000. Horseradish peroxidase– conjugated secondary antibodies and ECL reagents were used to detect primary antibodies. ImageJ 2 [96] was used to quantify band intensity. The relative intensity of the protein of interest was calculated by dividing the band intensity by the corresponding tubulin band intensity, then dividing by the band intensity in non-silenced control plant.

### RNA sequencing, differential expression analysis, and their functional enrichment, and predicting candidate PD genes

RNA was isolated from plant tissue from eighth/ninth leaves after 12-14 days of VIGS using Qiagen RNeasy® Plant Mini Kit and according to manufacturer protocol (Qiagen, Cat No. 74904) and digested with DNase I (Roche). RNA, approximately 25 to 500 ng/µl, was used for microfluidics analysis using a Bioanalyzer 2100 (Agilent Technologies, Santa Clara, CA) to assess RNA integrity using the Plant RNA Pico Assay chip following the manufacturer’s instructions. RNA Integrity Number (RIN) values were calculated to evaluate the quality of RNA samples and the RNA used for RNA seq analysis. RNA sequencing was performed by Novogene. Raw reads were analyzed for quality by FASTQC, and reads were mapped to the reference genome using HISAT2 [97]. The reads were mapped to the N. benthamiana V2.6.1 reference genome. The reference genome and gene model annotation files were all downloaded from the genome website (http://solgenomics.net/organism/Nicotiana_benthamiana/genome).

Gene counts were analyzed for gene-level differential expression using DESeq2 [98]. Significantly differentially expressed genes (DEGs) were identified with an adjusted P-value of less than 0.05. Heatmap of DEGs were made and clustered into six groups using k-means using iDEP 2.01 https://bioinformatics.sdstate.edu/idep/) web application [99]. More details are presented in the Supporting Information. For predicting candidate PD genes from DEGs, Arabidopsis orthologs of DEGs were obtained from The Arabidopsis Information Resource (TAIR) database. TAIR IDs were used for in silico prediction of PD genes using the PIP1 pipeline [59].

### Statistical analyses

Student’s t-test was used to determine statistical significance in the qPCR gene expression analysis, as well as to determine the levels of tetrapyrrole intermediates and quantify PD clusters. For movement assays, differences between mock and treatment (ALA, heme, or flg22) or between non-silenced and silenced plants were analyzed using the Bootstrap method [100] or Mann-Whitney U test in GraphPad Prism. p values under 0.05 were considered significant.

## ACKNOWLEDGEMENTS

We thank Dr. Barry Bruce (University of Tennessee, Knoxville) and members of the Burch-Smith lab for critical review of the manuscript.

## DATA AVAILABILITY

The RNA-seq data was deposited in the NCBI SRA under the accession number PRJNA1301871.

## REFERENCES

1. Paultre DS, Gustin MP, Molnar A, Oparka KJ. Lost in transit: long-distance trafficking and phloem unloading of protein signals in Arabidopsis homografts. Plant Cell. 2016. doi: 10.1105/tpc.16.00249. PubMed PMID: 27600534; PubMed Central PMCID: PMCPMC5059797.

2. Brunkard JO, Runkel AM, Zambryski PC. The cytosol must flow: intercellular transport through plasmodesmata. Current opinion in cell biology. 2015;35:13–20. doi: 10.1016/j.ceb.2015.03.003. PubMed PMID: 25847870.

3. Sager R, Lee JY. Plasmodesmata in integrated cell signalling: insights from development and environmental signals and stresses. J Exp Bot. 2014;65(22):6337–58. doi: 10.1093/jxb/eru365. PubMed PMID: 25262225; PubMed Central PMCID: PMC4303807.

4. Burch-Smith TM, Zambryski PC. Plasmodesmata paradigm shift: regulation from without versus within. Annu Rev Plant Biol. 2012;63:239–60. Epub 2011/12/06. doi: 10.1146/annurev-arplant-042811-105453. PubMed PMID: 22136566.

5. Kim I, Zambryski PC. Cell-to-cell communication via plasmodesmata during Arabidopsis embryogenesis. Curr Opin Plant Biol. 2005;8(6):593–9. Epub 2005/10/07. doi: S1369-5266(05)00136-6 [pii] 10.1016/j.pbi.2005.09.013. PubMed PMID: 16207533.

6. Burch-Smith TM, Stonebloom S, Xu M, Zambryski PC. Plasmodesmata during development: re-examination of the importance of primary, secondary, and branched plasmodesmata structure versus function. Protoplasma. 2011;248(1):61–74. Epub 2010/12/22. doi: 10.1007/s00709-010-0252-3. PubMed PMID: 21174132; PubMed Central PMCID: PMC3025111.

7. Reagan BC, Ganusova EE, Fernandez JC, McCray TN, Burch-Smith TM. RNA on the move: the plasmodesmata perspective. Plant Sci. 2018;In press.

8. Epel BL. Plant viruses spread by diffusion on ER-associated movement-protein-rafts through plasmodesmata gated by viral induced host beta-1,3-glucanases. Semin Cell Dev Biol. 2009;20(9):1074–81. Epub 2009/06/09. doi: S1084-9521(09)00109-8 [pii] 10.1016/j.semcdb.2009.05.010. PubMed PMID: 19501662.

9. Tee EE, Faulkner C. Plasmodesmata and intercellular molecular traffic control. The New phytologist. 2024;243(1):32–47. Epub 20240317. doi: 10.1111/nph.19666. PubMed PMID: 38494438.

10. Bayer EM, Benitez-Alfonso Y. Plasmodesmata: Channels Under Pressure. Annu Rev Plant Biol. 2024;75(1):291–317. Epub 20240702. doi: 10.1146/annurev-arplant-070623-093110. PubMed PMID: 38424063.

11. Zanini AA, Burch-Smith TM. New insights into plasmodesmata: complex ‘protoplasmic connecting threads’. J Exp Bot. 2024;75(18):5557–67. doi: 10.1093/jxb/erae307. PubMed PMID: 39001658; PubMed Central PMCID: PMCPMC11427835.

12. Burch-Smith TM, Brunkard JO, Choi YG, Zambryski PC. Organelle-nucleus cross-talk regulates plant intercellular communication via plasmodesmata. Proc Natl Acad Sci U S A. 2011;108(51):E1451-60. Epub 2011/11/23. doi: 1117226108 [pii] 10.1073/pnas.1117226108. PubMed PMID: 22106293; PubMed Central PMCID: PMC3251100.

13. Ehlers K, Kollmann R. Primary and secondary plasmodesmata: structure, origin, and functioning. Protoplasma. 2001;216(1-2):1–30. Epub 2001/12/06. PubMed PMID: 11732191.

14. Roberts IM, Boevink P, Roberts AG, Sauer N, Reichel C, Oparka KJ. Dynamic changes in the frequency and architecture of plasmodesmata during the sink-source transition in tobacco leaves. Protoplasma. 2001;218(1-2):31–44. PubMed PMID: 11732318.

15. Azim MF, Burch-Smith TM. Organelles-nucleus-plasmodesmata signaling (ONPS): an update on its roles in plant physiology, metabolism and stress responses. Curr Opin Plant Biol. 2020;58:48–59. Epub 2020/11/17. doi: 10.1016/j.pbi.2020.09.005. PubMed PMID: 33197746.

16. Kobayashi K, Otegui MS, Krishnakumar S, Mindrinos M, Zambryski P. INCREASED SIZE EXCLUSION LIMIT 2 encodes a putative DEVH box RNA helicase involved in plasmodesmata function during Arabidopsis embryogenesis. Plant Cell. 2007;19(6):1885–97. Epub 2007/07/03. doi: tpc.106.045666 [pii] 10.1105/tpc.106.045666. PubMed PMID: 17601829; PubMed Central PMCID: PMC1955720.

17. Bobik K, McCray TN, Ernest B, Fernandez JC, Howell KA, Lane T, et al. The chloroplast RNA helicase ISE2 is required for multiple chloroplast RNA processing steps in Arabidopsis thaliana. Plant J. 2017;91(1):114–31. Epub 2017/03/28. doi: 10.1111/tpj.13550. PubMed PMID: 28346704.

18. Carlotto N, Wirth S, Furman N, Ferreyra Solari N, Ariel F, Crespi M, et al. The chloroplastic DEVH-box RNA helicase INCREASED SIZE EXCLUSION LIMIT 2 involved in plasmodesmata regulation is required for group II intron splicing. Plant, cell & environment. 2016;39(1):165–73. doi: 10.1111/pce.12603. PubMed PMID: 26147377.

19. Schreier TB, Muller KH, Eicke S, Faulkner C, Zeeman SC, Hibberd JM. Plasmodesmal connectivity in C(4) Gynandropsis gynandra is induced by light and dependent on photosynthesis. The New phytologist. 2024;241(1):298–313. Epub 20231026. doi: 10.1111/nph.19343. PubMed PMID: 37882365.

20. Ganusova EE, Reagan BC, Fernandez JC, Azim MF, Sankoh AF, Freeman KM, et al. Chloroplast-to-nucleus retrograde signalling controls intercellular trafficking via plasmodesmata formation. Philos Trans R Soc Lond B Biol Sci. 2020;375(1801):20190408. Epub 2020/05/05. doi: 10.1098/rstb.2019.0408. PubMed PMID: 32362251; PubMed Central PMCID: PMCPMC7209952.

21. Bobik K, Fernandez JC, Hardin SR, Ernest B, Ganusova EE, Staton ME, et al. The essential chloroplast ribosomal protein uL15c interacts with the chloroplast RNA helicase ISE2 and affects intercellular trafficking through plasmodesmata. The New phytologist. 2019;221(2):850–65. doi: 10.1111/nph.15427. PubMed PMID: 30192000.

22. Surpin M, Larkin RM, Chory J. Signal transduction between the chloroplast and the nucleus. Plant Cell. 2002;14 Suppl:S327–38. PubMed PMID: 12045286.

23. Glasser C, Haberer G, Finkemeier I, Pfannschmidt T, Kleine T, Leister D, et al. Meta-analysis of retrograde signaling in Arabidopsis thaliana reveals a core module of genes embedded in complex cellular signaling networks. Mol Plant. 2014;7(7):1167–90. doi: 10.1093/mp/ssu042. PubMed PMID: 24719466.

24. Pogson BJ, Woo NS, Forster B, Small ID. Plastid signalling to the nucleus and beyond. Trends Plant Sci. 2008;13(11):602–9. Epub 2008/10/08. doi: S1360-1385(08)00227-6 [pii] 10.1016/j.tplants.2008.08.008. PubMed PMID: 18838332.

25. Gray JC, Sullivan JA, Wang JH, Jerome CA, MacLean D. Coordination of plastid and nuclear gene expression. Philos Trans R Soc Lond B Biol Sci. 2003;358(1429):135-44; discussion 44-5. Epub 2003/02/22. doi: 10.1098/rstb.2002.1180. PubMed PMID: 12594922; PubMed Central PMCID: PMCPMC1693108.

26. Susek RE, Ausubel FM, Chory J. Signal transduction mutants of Arabidopsis uncouple nuclear CAB and RBCS gene expression from chloroplast development. Cell. 1993;74(5):787–99. PubMed PMID: 7690685.

27. Woodson JD, Perez-Ruiz JM, Chory J. Heme synthesis by plastid ferrochelatase I regulates nuclear gene expression in plants. Curr Biol. 2011;21(10):897–903. PubMed PMID: 21565502.

28. Larkin RM, Alonso JM, Ecker JR, Chory J. GUN4, a regulator of chlorophyll synthesis and intracellular signaling. Science. 2003;299(5608):902-6. PubMed PMID: 12574634.

29. Mochizuki N, Brusslan JA, Larkin R, Nagatani A, Chory J. Arabidopsis genomes uncoupled 5 (GUN5) mutant reveals the involvement of Mg-chelatase H subunit in plastid-to-nucleus signal transduction. Proc Natl Acad Sci U S A. 2001;98(4):2053–8. PubMed PMID: 11172074.

30. Koussevitzky S, Nott A, Mockler TC, Hong F, Sachetto-Martins G, Surpin M, et al. Signals from chloroplasts converge to regulate nuclear gene expression. Science. 2007;316(5825):715-9. PubMed PMID: 17395793.

31. He RR, Li YZ, Bernards MA, Wang AM. Turnip mosaic virus selectively subverts a PR-5 thaumatin-like, plasmodesmal protein to promote viral infection. New Phytologist. 2025;245(1):299–317. doi: 10.1111/nph.20233. PubMed PMID: WOS:001356959600001.

32. Stephenson PG, Terry MJ. Light signalling pathways regulating the Mg-chelatase branchpoint of chlorophyll synthesis during de-etiolation in Arabidopsis thaliana. Photochem Photobiol Sci. 2008;7(10):1243–52. Epub 20080723. doi: 10.1039/b802596g. PubMed PMID: 18846290.

33. Strand A, Asami T, Alonso J, Ecker JR, Chory J. Chloroplast to nucleus communication triggered by accumulation of Mg-protoporphyrinIX. Nature. 2003;421(6918):79-83. PubMed PMID: 12511958.

34. Voigt C, Oster U, Bornke F, Jahns P, Dietz KJ, Leister D, et al. In-depth analysis of the distinctive effects of norflurazon implies that tetrapyrrole biosynthesis, organellar gene expression and ABA cooperate in the GUN-type of plastid signalling. Physiol Plant. 2010;138(4):503–19. PubMed PMID: 20028479.

35. Moulin M, McCormac AC, Terry MJ, Smith AG. Tetrapyrrole profiling in Arabidopsis seedlings reveals that retrograde plastid nuclear signaling is not due to Mg-protoporphyrin IX accumulation. Proc Natl Acad Sci U S A. 2008;105(39):15178–83. PubMed PMID: 18818314.

36. von Gromoff ED, Alawady A, Meinecke L, Grimm B, Beck CF. Heme, a plastid-derived regulator of nuclear gene expression in Chlamydomonas. Plant Cell. 2008;20(3):552–67. PubMed PMID: 18364467.

37. Zhang L, Hach A. Molecular mechanism of heme signaling in yeast: the transcriptional activator Hap1 serves as the key mediator. Cell Mol Life Sci. 1999;56(5-6):415–26. PubMed PMID: 11212295.

38. Chen C, Samuel TK, Sinclair J, Dailey HA, Hamza I. An intercellular heme-trafficking protein delivers maternal heme to the embryo during development in C. elegans. Cell. 2011;145(5):720–31. doi: 10.1016/j.cell.2011.04.025. PubMed PMID: 21620137; PubMed Central PMCID: PMCPMC3104245.

39. Yin L, Wu N, Curtin JC, Qatanani M, Szwergold NR, Reid RA, et al. Rev-erbalpha, a heme sensor that coordinates metabolic and circadian pathways. Science. 2007;318(5857):1786-9. Epub 2007/11/17. doi: 10.1126/science.1150179. PubMed PMID: 18006707.

40. Chen Y, Nishimura K, Tokizawa M, Yamamoto YY, Oka Y, Matsushita T, et al. Alternative localization of HEME OXYGENASE 1 in plant cells regulates cytosolic heme catabolism. Plant Physiol. 2024;195(4):2937–51. doi: 10.1093/plphys/kiae288. PubMed PMID: 38805221.

41. Page MT, Garcia-Becerra T, Smith AG, Terry MJ. Overexpression of chloroplast-targeted ferrochelatase 1 results in a genomes uncoupled chloroplast-to-nucleus retrograde signalling phenotype. Philos Trans R Soc Lond B Biol Sci. 2020;375(1801):20190401. Epub 20200504. doi: 10.1098/rstb.2019.0401. PubMed PMID: 32362255; PubMed Central PMCID: PMCPMC7209946.

42. Espinas NA, Kobayashi K, Sato Y, Mochizuki N, Takahashi K, Tanaka R, et al. Allocation of Heme Is Differentially Regulated by Ferrochelatase Isoforms in Arabidopsis Cells. Frontiers in plant science. 2016;7:1326. Epub 2016/09/16. doi: 10.3389/fpls.2016.01326. PubMed PMID: 27630653; PubMed Central PMCID: PMCPMC5005420.

43. Benitez M, Hernandez-Hernandez V, Newman SA, Niklas KJ. Dynamical Patterning Modules, Biogeneric Materials, and the Evolution of Multicellular Plants. Frontiers in plant science. 2018;9:871. Epub 2018/08/01. doi: 10.3389/fpls.2018.00871. PubMed PMID: 30061903; PubMed Central PMCID: PMCPMC6055014.

44. Brunkard JO, Zambryski PC. Plasmodesmata enable multicellularity: new insights into their evolution, biogenesis, and functions in development and immunity. Curr Opin Plant Biol. 2017;35:76–83. doi: 10.1016/j.pbi.2016.11.007. PubMed PMID: 27889635.

45. Tanaka R, Kobayashi K, Masuda T. Tetrapyrrole Metabolism in *Arabidopsis thaliana*. The Arabidopsis Book. 2011;2011(9).

46. Burch-Smith TM, Zambryski PC. Loss of INCREASED SIZE EXCLUSION LIMIT (ISE)1 or ISE2 increases the formation of secondary plasmodesmata. Curr Biol. 2010;20(11):989–93. Epub 2010/05/04. doi: S0960-9822(10)00439-2 [pii] 10.1016/j.cub.2010.03.064. PubMed PMID: 20434343; PubMed Central PMCID: PMC2902234.

47. Mochizuki N, Tanaka R, Grimm B, Masuda T, Moulin M, Smith AG, et al. The cell biology of tetrapyrroles: a life and death struggle. Trends Plant Sci. 2010;15(9):488–98. PubMed PMID: 20598625.

48. Shimizu T, Yasuda R, Mukai Y, Tanoue R, Shimada T, Imamura S, et al. Proteomic analysis of haem-binding protein from Arabidopsis thaliana and Cyanidioschyzon merolae. Philos Trans R Soc Lond B Biol Sci. 2020;375(1801):20190488. Epub 20200504. doi: 10.1098/rstb.2019.0488. PubMed PMID: 32362261; PubMed Central PMCID: PMCPMC7209954.

49. Papenbrock J, Mock HP, Tanaka R, Kruse E, Grimm B. Role of magnesium chelatase activity in the early steps of the tetrapyrrole biosynthetic pathway. Plant Physiol. 2000;122(4):1161–9. PubMed PMID: 10759511.

50. Scharfenberg M, Mittermayr L, E VONR-L, Schlicke H, Grimm B, Leister D, et al. Functional characterization of the two ferrochelatases in Arabidopsis thaliana. Plant, cell & environment. 2015;38(2):280–98. Epub 2013/12/18. doi: 10.1111/pce.12248. PubMed PMID: 24329537.

51. Woodson JD, Joens MS, Sinson AB, Gilkerson J, Salome PA, Weigel D, et al. Ubiquitin facilitates a quality-control pathway that removes damaged chloroplasts. Science. 2015;350(6259):450-4. Epub 2015/10/24. doi: 10.1126/science.aac7444. PubMed PMID: 26494759.

52. Papenbrock J, Mishra S, Mock HP, Kruse E, Schmidt EK, Petersmann A, et al. Impaired expression of the plastidic ferrochelatase by antisense RNA synthesis leads to a necrotic phenotype of transformed tobacco plants. Plant J. 2001;28(1):41–50. PubMed PMID: 11696185.

53. Richter AS, Banse C, Grimm B. The GluTR-binding protein is the heme-binding factor for feedback control of glutamyl-tRNA reductase. eLife. 2019;8. Epub 20190613. doi: 10.7554/eLife.46300. PubMed PMID: 31194674; PubMed Central PMCID: PMCPMC6597238.

54. Kumar S, Bandyopadhyay U. Free heme toxicity and its detoxification systems in human. Toxicol Lett. 2005;157(3):175–88. Epub 20050407. doi: 10.1016/j.toxlet.2005.03.004. PubMed PMID: 15917143.

55. Wu Y, Li J, Wang J, Dawuda MM, Liao W, Meng X, et al. Heme is involved in the exogenous ALA-promoted growth and antioxidant defense system of cucumber seedlings under salt stress. BMC plant biology. 2022;22(1):329. Epub 20220708. doi: 10.1186/s12870-022-03717-3. PubMed PMID: 35804328; PubMed Central PMCID: PMCPMC9264505.

56. Brunkard JO, Zambryski P. Plant Cell-Cell Transport via Plasmodesmata Is Regulated by Light and the Circadian Clock. Plant Physiol. 2019;181(4):1459–67. Epub 2019/10/12. doi: 10.1104/pp.19.00460. PubMed PMID: 31601643; PubMed Central PMCID: PMCPMC6878007.

57. Epel BL, Erlanger MA. Light regulates symplastic communication in etiolated corn seedlings. Physiologia Plantarum. 1991;83:149–53.

58. Jimenez-Gongora T, Kim SK, Lozano-Duran R, Zipfel C. Flg22-Triggered Immunity Negatively Regulates Key BR Biosynthetic Genes. Frontiers in plant science. 2015;6:981. Epub 20151109. doi: 10.3389/fpls.2015.00981. PubMed PMID: 26617621; PubMed Central PMCID: PMCPMC4637422.

59. Kirk P, Amsbury S, German L, Gaudioso-Pedraza R, Benitez-Alfonso Y. A comparative meta-proteomic pipeline for the identification of plasmodesmata proteins and regulatory conditions in diverse plant species. BMC biology. 2022;20(1):128. Epub 20220602. doi: 10.1186/s12915-022-01331-1. PubMed PMID: 35655273; PubMed Central PMCID: PMCPMC9164936.

60. Raffaele S, Bayer E, Lafarge D, Cluzet S, German Retana S, Boubekeur T, et al. Remorin, a solanaceae protein resident in membrane rafts and plasmodesmata, impairs potato virus X movement. Plant Cell. 2009;21(5):1541–55. Epub 2009/05/28. doi: tpc.108.064279 [pii] 10.1105/tpc.108.064279. PubMed PMID: 19470590; PubMed Central PMCID: PMC2700541.

61. Alazem M, Nuzzi SP, Burch-Smith TM. Viral Movement Proteins and Plasmodesmata: turning gatekeepers into gateways. J Exp Bot. 2025:Accepted.

62. Saheki Y, De Camilli P. Endoplasmic Reticulum-Plasma Membrane Contact Sites. Annual Review of Biochemistry, Vol 86. 2017;86:659–84. doi: 10.1146/annurev-biochem-061516-044932. PubMed PMID: WOS:000407725800027.

63. Dharan R, Sorkin R. Tetraspanin proteins in membrane remodeling processes. Journal of cell science. 2024;137(14). Epub 20240725. doi: 10.1242/jcs.261532. PubMed PMID: 39051897; PubMed Central PMCID: PMCPMC7617763.

64. Boavida LC, Qin P, Broz M, Becker JD, McCormick S. Arabidopsis tetraspanins are confined to discrete expression domains and cell types in reproductive tissues and form homo- and heterodimers when expressed in yeast. Plant Physiol. 2013;163(2):696–712. Epub 20130814. doi: 10.1104/pp.113.216598. PubMed PMID: 23946353; PubMed Central PMCID: PMCPMC3793051.

65. Johnston MG, Breakspear A, Samwald S, Zhang D, Papp D, Faulkner C, et al. Comparative phyloproteomics identifies conserved plasmodesmal proteins. J Exp Bot. 2023;74(6):1821–35. doi: 10.1093/jxb/erad022. PubMed PMID: 36639877; PubMed Central PMCID: PMCPMC10049917.

66. Stahl Y, Grabowski S, Bleckmann A, Kuhnemuth R, Weidtkamp-Peters S, Pinto KG, et al. Moderation of Arabidopsis root stemness by CLAVATA1 and ARABIDOPSIS CRINKLY4 receptor kinase complexes. Curr Biol. 2013;23(5):362–71. Epub 20130207. doi: 10.1016/j.cub.2013.01.045. PubMed PMID: 23394827.

67. Fitzgibbon J, Beck M, Zhou J, Faulkner C, Robatzek S, Oparka K. A developmental framework for complex plasmodesmata formation revealed by large-scale imaging of the Arabidopsis leaf epidermis. Plant Cell. 2013;25(1):57–70. doi: 10.1105/tpc.112.105890. PubMed PMID: 23371949; PubMed Central PMCID: PMC3584549.

68. Sankoh AF, Adjei J, Roberts D, Burch-Smith TM. Comparing methods for detection and quantification of plasmodesmal callose in Nicotiana benthamiana leaves during defense responses. Mol Plant Microbe Interact. 2024. Epub 20240220. doi: 10.1094/MPMI-09-23-0152-SC. PubMed PMID: 38377039.

69. Koreny L, Obornik M, Horakova E, Waller RF, Lukes J. The convoluted history of haem biosynthesis. Biol Rev Camb Philos Soc. 2022;97(1):141–62. Epub 20210902. doi: 10.1111/brv.12794. PubMed PMID: 34472688.

70. Layer G, Reichelt J, Jahn D, Heinz DW. Structure and function of enzymes in heme biosynthesis. Protein Science. 2010;19(6):1137–61. doi: 10.1002/pro.405.

71. Suzuki T, Masuda T, Singh DP, Tan FC, Tsuchiya T, Shimada H, et al. Two types of ferrochelatase in photosynthetic and nonphotosynthetic tissues of cucumber: their difference in phylogeny, gene expression, and localization. J Biol Chem. 2002;277(7):4731–7. PubMed PMID: 11675381.

72. Chow KS, Singh DP, Walker AR, Smith AG. Two different genes encode ferrochelatase in Arabidopsis: mapping, expression and subcellular targeting of the precursor proteins. Plant J. 1998;15(4):531–41. PubMed PMID: 9753778.

73. von Gromoff ED, Schroda M, Oster U, Beck CF. Identification of a plastid response element that acts as an enhancer within the Chlamydomonas HSP70A promoter. Nucleic Acids Res. 2006;34(17):4767–79. PubMed PMID: 16971458.

74. Shan Y, Lambrecht RW, Ghaziani T, Donohue SE, Bonkovsky HL. Role of Bach-1 in regulation of heme oxygenase-1 in human liver cells: insights from studies with small interfering RNAS. J Biol Chem. 2004;279(50):51769–74. Epub 20041001. doi: 10.1074/jbc.M409463200. PubMed PMID: 15465821.

75. Sun J, Brand M, Zenke Y, Tashiro S, Groudine M, Igarashi K. Heme regulates the dynamic exchange of Bach1 and NF-E2-related factors in the Maf transcription factor network. Proc Natl Acad Sci U S A. 2004;101(6):1461–6. Epub 20040127. doi: 10.1073/pnas.0308083100. PubMed PMID: 14747657; PubMed Central PMCID: PMCPMC341742.

76. Zitomer RS, Lowry CV. Regulation of gene expression by oxygen in Saccharomyces cerevisiae. Microbiological reviews. 1992;56(1):1–11. doi: 10.1128/mr.56.1.1-11.1992. PubMed PMID: 1579104; PubMed Central PMCID: PMCPMC372851.

77. Guarente L, Mason T. Heme regulates transcription of the CYC1 gene of S. cerevisiae via an upstream activation site. Cell. 1983;32(4):1279–86. doi: 10.1016/0092-8674(83)90309-4. PubMed PMID: 6301690.

78. Joshi B, Morley SJ, Rhoads RE, Pain VM. Inhibition of Protein Synthesis by the Heme-Controlled Eif-2α kinase Leads to the Appearance of mRNA-Containing 48S Complexes that Contain eIF-4E but Lack Methionyl-tRNAf. European Journal of Biochemistry. 1995;228(1):31–8. doi: 10.1111/j.1432-1033.1995.0031o.x.

79. Chen JJ, Yang JM, Petryshyn R, Kosower N, London IM. Disulfide bond formation in the regulation of eIF-2 alpha kinase by heme. J Biol Chem. 1989;264(16):9559–64. PubMed PMID: 2722851.

80. Tang XD, Xu R, Reynolds MF, Garcia ML, Heinemann SH, Hoshi T. Haem can bind to and inhibit mammalian calcium-dependent Slo1 BK channels. Nature. 2003;425(6957):531-5. doi: 10.1038/nature02003. PubMed PMID: 14523450.

81. Zhao C, Wang Y, Chan KX, Marchant DB, Franks PJ, Randall D, et al. Evolution of chloroplast retrograde signaling facilitates green plant adaptation to land. Proc Natl Acad Sci U S A. 2019;116(11):5015–20. Epub 2019/02/26. doi: 10.1073/pnas.1812092116. PubMed PMID: 30804180; PubMed Central PMCID: PMCPMC6421419.

82. Wen B, Grimm B. A genetically encoded fluorescent heme sensor detects free heme in plants. Plant Physiol. 2024;196(2):830–41. doi: 10.1093/plphys/kiae291. PubMed PMID: 38762898; PubMed Central PMCID: PMCPMC11444292.

83. Terry MJ, Smith AG. A model for tetrapyrrole synthesis as the primary mechanism for plastid-to-nucleus signaling during chloroplast biogenesis. Frontiers in plant science. 2013;4:14. Epub 2013/02/15. doi: 10.3389/fpls.2013.00014. PubMed PMID: 23407626; PubMed Central PMCID: PMCPMC3570980.

84. Thomas J, Weinstein JD. Measurement of Heme Efflux and Heme Content in Isolated Developing Chloroplasts 1. Plant Physiol. 1990;94(3):1414–23. doi: 10.1104/pp.94.3.1414.

85. Espinas NA, Kobayashi K, Takahashi S, Mochizuki N, Masuda T. Evaluation of unbound free heme in plant cells by differential acetone extraction. Plant Cell Physiol. 2012;53(7):1344–54. Epub 20120502. doi: 10.1093/pcp/pcs067. PubMed PMID: 22555813.

86. Fan T, Roling L, Meiers A, Brings L, Ortega-Rodes P, Hedtke B, et al. Complementation studies of the Arabidopsis fc1 mutant substantiate essential functions of ferrochelatase 1 during embryogenesis and salt stress. Plant, cell & environment. 2019;42(2):618–32. Epub 20181119. doi: 10.1111/pce.13448. PubMed PMID: 30242849.

87. Tanaka R, Tanaka A. Tetrapyrrole biosynthesis in higher plants. Annu Rev Plant Biol. 2007;58:321–46. PubMed PMID: 17227226.

88. Stonebloom S, Burch-Smith T, Kim I, Meinke D, Mindrinos M, Zambryski P. Loss of the plant DEAD-box protein ISE1 leads to defective mitochondria and increased cell-to-cell transport via plasmodesmata. Proc Natl Acad Sci U S A. 2009;106(40):17229–34. Epub 2009/10/07. doi: 0909229106 [pii] 10.1073/pnas.0909229106. PubMed PMID: 19805190; PubMed Central PMCID: PMC2761335.

89. Dmitrieva VA, Domashkina VV, Ivanova AN, Sukhov VS, Tyutereva EV, Voitsekhovskaja OV. Regulation of plasmodesmata in Arabidopsis leaves: ATP, NADPH and chlorophyll b levels matter. J Exp Bot. 2021;72(15):5534–52. doi: 10.1093/jxb/erab205. PubMed PMID: 33974689.

90. Brunkard JO, Runkel AM, Zambryski PC. Chloroplasts extend stromules independently and in response to internal redox signals. Proc Natl Acad Sci U S A. 2015;112(32):10044–9. doi: 10.1073/pnas.1511570112. PubMed PMID: 26150490; PubMed Central PMCID: PMC4538653.

91. Liu Y, Schiff M, Marathe R, Dinesh-Kumar SP. Tobacco Rar1, EDS1 and NPR1/NIM1 like genes are required for N-mediated resistance to tobacco mosaic virus. Plant J. 2002;30(4):415–29. Epub 2002/05/25. doi: 1297 [pii]. PubMed PMID: 12028572.

92. Fernandez JC, Burch-Smith TM. Investigating Plasmodesmata Function in Arabidopsis Thaliana Using a Low-Pressure Bombardment System and GFP Movement Assay. Methods Mol Biol. 2022;2457:273–83. doi: 10.1007/978-1-0716-2132-5_18. PubMed PMID: 35349147.

93. Cheval C, Samwald S, Johnston MG, de Keijzer J, Breakspear A, Liu X, et al. Chitin perception in plasmodesmata characterizes submembrane immune-signaling specificity in plants. Proc Natl Acad Sci U S A. 2020;117(17):9621–9. Epub 2020/04/15. doi: 10.1073/pnas.1907799117. PubMed PMID: 32284410; PubMed Central PMCID: PMCPMC7196898.

94. Ishikawa K, Tamura K, Fukao Y, Shimada T. Structural and functional relationships between plasmodesmata and plant endoplasmic reticulum-plasma membrane contact sites consisting of three synaptotagmins. The New phytologist. 2020;226(3):798–808. Epub 20200131. doi: 10.1111/nph.16391. PubMed PMID: 31869440.

95. Klinkenberg J. Extraction of Chloroplast Proteins from Transiently Transformed Nicotiana benthamiana Leaves. Bio-protocol. 2014;4(18):e1238. doi: 10.21769/BioProtoc.1238.

96. Schneider CA, Rasband WS, Eliceiri KW. NIH Image to ImageJ: 25 years of image analysis. Nature methods. 2012;9(7):671–5. PubMed PMID: 22930834.

97. Kim D, Langmead B, Salzberg SL. HISAT: a fast spliced aligner with low memory requirements. Nature methods. 2015;12(4):357–60. Epub 20150309. doi: 10.1038/nmeth.3317. PubMed PMID: 25751142; PubMed Central PMCID: PMCPMC4655817.

98. Love MI, Huber W, Anders S. Moderated estimation of fold change and dispersion for RNA-seq data with DESeq2. Genome Biol. 2014;15(12):550. doi: 10.1186/s13059-014-0550-8. PubMed PMID: 25516281; PubMed Central PMCID: PMCPMC4302049.

99. Ge SX, Son EW, Yao R. iDEP: an integrated web application for differential expression and pathway analysis of RNA-Seq data. BMC Bioinformatics. 2018;19(1):534. doi: 10.1186/s12859-018-2486-6.

100. Johnston MG, Faulkner C. A bootstrap approach is a superior statistical method for the comparison of non-normal data with differing variances. The New phytologist. 2021;230(1):23–6. Epub 20210129. doi: 10.1111/nph.17159. PubMed PMID: 33349922.

