## Supporting Information for "Heme acts a chloroplast-to-nucleus retrograde signal to regulate intercellular trafficking via plasmodesmata"

Supporting Figure S1

```
AtGUN4_AT3G59400          MAT--TNSLHHHHHSSPSYTHHRNNLHCQSHFGPTSLSLKQPTSAATFSLICSASSTSSS        58
Nbe.v1.1.chr09g39610      MATNSFKSIHHHHTI----IRRRHSIDC--PFSSSSFFLKPII--------KHPSSTSYS        46
Nbe.v1.1.chr16g37380      -------------------------MQE--NFGVDDD-------------------FYD         13
                                                   :.    *.  .                       .

AtGUN4_AT3G59400          T--TAVSAVSTTNASATTAETATIFDVLENHLVNQNFRQADEETRRLLIQISGEAAVKRG        116
Nbe.v1.1.chr09g39610      NHNLITFSLSPTSPATTPITPTTPFDILEQYLSAQDFRQADEETRRLLIVLAGEAAVKRG        106
Nbe.v1.1.chr16g37380      MLDDIQLSDSL---------------EIPLPTPAQDFRQADEETRRLLIVLAGEAAVKRG        58
                                 : *                 :      *:************* ::********

AtGUN4_AT3G59400          YVFFSEVKTISPEDLQAIDNLWIKHSDGRFGYSVQRKIWLKVKKDFTRFFVKVEWMKLLD        176
Nbe.v1.1.chr09g39610      YVFFSEVQFIQESDLKEIDSLWKKYSDNKFGYSVQKKIWNKVNRDFTSFFIKVGWMKKLE        166
Nbe.v1.1.chr16g37380      YVFFSEVQFIQESDLKEIDSLWKKYSDNKFGYSVQKKIWNKVNRDFTSYFIKVGWMKKLE        118
                          *******: *. .**: **.** *:**.:******:*** **::*** :*:** *** *:

AtGUN4_AT3G59400          -TEVVQYNYRAFPDEFKWELNDETPLGHLPLTNALRGTQLLKCVLSHPAFATADDNSGET        235
Nbe.v1.1.chr09g39610      SSEVEQYNYRAFPNEFIWELNEETPEGHLPLTNALRGTQLLSSIFTHPAFVEQGEEEKDE        226
Nbe.v1.1.chr16g37380      SSEVDQYNYRAFPNEFIWELNEETPDGHLPLTNALRGTQLLSSIFTHPALVEEGEEEKEE        178
                           :** ********:** ****:*** ***************..:::***:.  .::. :

AtGUN4_AT3G59400          EDELNR-GV---AVAKEQ-AGVGADKRVFKTNYSF  265
Nbe.v1.1.chr09g39610      EKQASFSGEDKSSVNKGGLLGGLRSKLFGKPDYSF  261
Nbe.v1.1.chr16g37380      EKQGSISGEDKRSVKKGGLLGGLRSKLFGKPDYSF  213
                          *.: .  *    :* *    *   .* . * :***
```

**Figure S1**. Multiple sequence alignment of *N. benthamiana* GUN4 homeologous proteins with *A. thaliana* (AtGUN4). Sequences of Nbe.v1.1.chr09g39610 and Nbe.v1.1.chr16g37380, were retrieved from the NbenBase DB [1] using the AtGUN4 sequence as a query. The conserved GUN4 domain was identified by the NCBI Conserved Domain Database (CDD) [2]. The GUN4 domain is highlighted with gray for AtGUN4, yellow for Nbe.v1.1.chr09g39610, and blue for Nbe.v1.1.chr16g37380.

Supporting Figure S2

```
AtGUN5_AT5G13630        MASLVYSPFTLSTSKAEHLSSLTNST--KHSFLRKKHRSTKPAKSFFKVKSAVSGNGLFT        58
Nbe.v1.1.chr04g34700    MASLVSSPFTLPNSKVEHLSSISQKHYFLHSFLPKKTNPT-YSKSPKKFQCNAIGNGLFT        59
Nbe.v1.1.chr14g15590    MASLVSSPFTLPNSKVEHLSSISQKHYFLHSFLPKKTNPT-YSKSPKKFQCNAIGNGLFT        59
                        ***** ***** .**.*****:::.    **** ** . *   :**  *.:. . ******

AtGUN5_AT5G13630        QTNPEVRRIVPIKRDNVPTVKIVYVVLEAQYQSSLSEAVQSLNKTSRFASYEVVGYLVEE        118
Nbe.v1.1.chr04g34700    QTTQELRRIVPENIQGLATVKIVYVVLEAQYQASLTAAVQTLNKNGKFASFEVVGYLVEE        119
Nbe.v1.1.chr14g15590    QTTQEVRRIVPENTQGLATVKIVYVVLEAQYQASLTAAVQTLNKNGKFASFEVVGYLVEE        119
                        **. *:***** : :.: **************:**: ***:***..:***:*********

AtGUN5_AT5G13630        LRDKNTYNNFCEDLKDANIFIGSLIFVEELAIKVKDAVEKERDRMDAVLVFPSMPEVMRL        178
Nbe.v1.1.chr04g34700    LRDENNYKMFCKDLEDANVFIGSLIFVEELALKVKTAVEKERDRLDAVLVFPSMPEVMRL        179
Nbe.v1.1.chr14g15590    LRDENTYKMLCKDLEDANVFIGSLIFVEELALKVKSAVEKERDRLDAVLVFPSMPEVMRL        179
                        ***:*.*: :*:**:***:************:*** ********:***************

AtGUN5_AT5G13630        NKLGSFSMSQLGQSKSPFFQLFKRKKQGSAGFADSMLKLVRTLPKVLKYLPSDKAQDARL        238
Nbe.v1.1.chr04g34700    NKLGSFSMSQLGQSKSPFFELFKKKKSSSAGFSDQMLKLVRTLPKVLKYLPSDKAQDARL        239
Nbe.v1.1.chr14g15590    NKLGSFSMSQLGQSKSPFFELFKKKKSSSAGFSDQMLKLVRTLPKVLKYLPSDKAQDARL        239
                        *******************:***:**..****:*.*************************

AtGUN5_AT5G13630        YILSLQFWLGGSPDNLQNFVKMISGSYVPALKGVKIEYSDPVLFLDTGIWHPLAPTMYDD        298
Nbe.v1.1.chr04g34700    YILSLQFWLGGSPDNLVNFLKMISGSYVPALKGMKIDYSDPVLYLDSGIWHPLAPCMYDD        299
Nbe.v1.1.chr14g15590    YILSLQFWLGGSPDNLVNFLKMISGSYVPALKGMKIDYSDPVLYLDSGIWHPLAPCMYDD        299
                        **************** **:*************:**:******:**:******** ****

AtGUN5_AT5G13630        VKEYWNWYDTRRDTNDSLKRKDATVVGLVLQRSHIVTGDDSHYVAVIMELEARGAKVVPI        358
Nbe.v1.1.chr04g34700    VKEYLNWYATRRDTNEKLKNSSAPVVGLVLQRSHIVTGDESHYVAVIMELEAKGAKVIPI        359
Nbe.v1.1.chr14g15590    VKEYLNWYATRRDTNEKLKSSNAPVVGLVLQRSHIVTGDESHYVAVIMELEAKGAKVIPI        359
                        **** *** ******..**  ..* ***************:************:****:**

AtGUN5_AT5G13630        FAGGLDFSGPVEKYFVDPVSKQPIVNSAVSLTGFALVGGPARQDHPRAIEALKKLDVPYL        418
Nbe.v1.1.chr04g34700    FAGGLDFSGPVERYFIDPITKKPFVNSVISLTGFALVGGPARQDHPRAIEALMKLDVPYI        419
Nbe.v1.1.chr14g15590    FAGGLDFSGPVERYFIDPITKKPFVNSVISLTGFALVGGPARQDHPRAIEALMKLDVPYI        419
                        ***********:**:**::*:*:***.:*********************** ******:

AtGUN5_AT5G13630        VAVPLVFQTTEEWLNSTLGLHPIQVALQVALPELDGAMEPIVFAGRDPRTGKSHALHKRV        478
Nbe.v1.1.chr04g34700    VALPLVFQTTEEWLNSTLGLHPIQVALQVALPELDGGMEPIVFAGRDPRTGKSHALHKRV        479
Nbe.v1.1.chr14g15590    VALPLVFQTTEEWLNSTLGLHPIQVALQVALPELDGGMEPIVFSGRDPRTGKSHALHKRV        479
                        **:*********************************.******:****************

AtGUN5_AT5G13630        EQLCIRAIRWGELKRKTKAEKKLAITVFSFPPDKGNVGTAAYLNVFASIFSVLRDLKRDG        538
Nbe.v1.1.chr04g34700    EQLCTRAIKWGELKRKTKAEKRLAITVFSFPPDKGNVGTAAYLNVFASIYSVLKELKKDG        539
Nbe.v1.1.chr14g15590    EQLCTRAIKWGELKRKTKAEKRLAITVFSFPPDKGNVGTAAYLNVFASIYSVLKDLKKDG        539
                        **** ***:************:****************************:***::**:**

AtGUN5_AT5G13630        YNVEGLPENAETLIEEIIHDKEAQFSSPNLNVAYKMGVREYQDLTPYANALEENWGKPPG        598
Nbe.v1.1.chr04g34700    YNIEGLPETSAQMIEEVIHDKEAQFSSPNLNIAYKMNVREYQKLTPYATALEENWGKAPG        599
Nbe.v1.1.chr14g15590    YNVEGLPDTSAELIEEVIHDKEAQFSSPNLNIAYKMSVREYQKLTPYATALEENWGKAPG        599
                        **:*****::.:  :***:**************:****.*****.*****.******** **

AtGUN5_AT5G13630        NLNSDGENLLVYGKAYGNVFIGVQPTFGYEGDPMRLLFSKSASPHHGFAAYYSYVEKIFK        658
Nbe.v1.1.chr04g34700    NLNSDGENLLVYGKQYGNVFIGVQPTFGYEGDPMRLLFSKSASPHHGFAAYYSFVEKIFK        659
Nbe.v1.1.chr14g15590    NLNSDGENLLVYGKQYGNVFIGVQPTFGYEGDPMRLLFSKSASPHHGFAAYYSFVEKIFK        659
                        ************** **************************************:******

AtGUN5_AT5G13630        ADAVLHFGTHGSLEFMPGKQVGMSDACFPDSLIGNIPNVYYYAANNPSEATIAKRRSYAN        718
Nbe.v1.1.chr04g34700    ADAVLHFGTHGSLEFMPGKQVGMSDACFPDSLIGNIPNVYYYAANNPSEATIAKRRSYAN        719
Nbe.v1.1.chr14g15590    ADAVLHFGTHGSLEFMPGKQVGMSDACFPDSLIGNIPNVYYYAANNPSEATIAKRRSYAN        719
                        ************************************************************

AtGUN5_AT5G13630        TISYLTPPAENAGLYKGLKQLSELISSYQSLKDTGRGPQIVSSIISTAKQCNLDKDVDLP        778
Nbe.v1.1.chr04g34700    TISYLTPPAENAGLYKGLKQLSELISSYQSLKDSGRGQQIVNSIISTARQCNLDKDVDLP        779
Nbe.v1.1.chr14g15590    TISYLTPPAENAGLYKGLKQLSELISSYQSLKDSGRGQQIVNSIISTARQCNLDKDVDLP        779
                        *********************************:*** ***.******:***********
```

```
AtGUN5_AT5G13630          DEGLELSPKDRDSVVGKVYSKIMEIESRLLPCGLHVIGEPPSAMEAVATLVNIAALDRPE        838
Nbe.v1.1.chr04g34700      EEGEEISAKERDLVVGKVYSKIMEIESRLLPCGLHIIGEPPTAMEAVATLVNIAALDRPE        839
Nbe.v1.1.chr14g15590      EEGEEISAKERDLVVGKVYSKIMEIESRLLPCGLHIIGEPPTAMEAVATLVNIAALDRPE        839
                          :** *:* *:** **********************:*****:******************

AtGUN5_AT5G13630          DEISALPSILAECVGREIEDVYRGSDKGILSDVELLKEITDASRGAVSAFVEKTTNSKGQ        898
Nbe.v1.1.chr04g34700      DGISALPSILAATVGRSIEEIYRGNDQGILRDVELLRQITEASRGAISAFVERTTNNKGQ        899
Nbe.v1.1.chr14g15590      EGISALPSILAATVGRSIEEIYRGNDQGILRDVELLRQITEASRGAISAFVERTTNNKGQ        899
                          : *********  ***.**::***.*:*** *****::**:*****:*****:***.***

AtGUN5_AT5G13630          VVDVSDKLTSLLGFGINEPWVEYLSNTKFYRANRDKLRTVFGFLGECLKLVVMDNELGSL        958
Nbe.v1.1.chr04g34700      VVNVNDKLTSILGFGINEPWVQYLSNTQFYRADREKLRVLFQFLGECLKLIVANNEVGSL        959
Nbe.v1.1.chr14g15590      VVNVNDKLTSILGFGINEPWIQYLSNTQFYRADREKLRVLFQFLGECLKLIVANNEVGSL        959
                          **:*.*****:*********::*****:****:*:***.:* ********:* :**:***

AtGUN5_AT5G13630          MQALEGKYVEPGPGGDPIRNPKVLPTGKNIHALDPQAIPTTAAMASAKIVVERLVERQKL        1018
Nbe.v1.1.chr04g34700      KQALEGKYVEPGPGGDPIRNPKVLPTGKNIHALDPQAIPTVAAVQSAKIVVERLLERQKA        1019
Nbe.v1.1.chr14g15590      KQALEGKYVEPGPGGDPIRNPKVLPTGKNIHALDPQAIPTVAAVQSAKVVVERLLERQKA        1019
                           ***************************************.**: ***:*****:****

AtGUN5_AT5G13630          ENEGKYPETIALVLWGTDNIKTYGESLGQVLWMIGVRPIADTFGRVNRVEPVSLEELGRP        1078
Nbe.v1.1.chr04g34700      DNGGKYPETVALVLWGTDNIKTYGESLAQVMWMIGVRPVTDTLGRVNQVEPVSLEELGRP        1079
Nbe.v1.1.chr14g15590      DNGGKYPETVALVLWGTDNIKTYGESLAQVMWMIGVRPVTDSLGRVNRVEPVSLEELGRP        1079
                          :* ******:*****************.**:*******::*::****:************

AtGUN5_AT5G13630          RIDVVVNCSGVFRDLFINQMNLLDRAIKMVAELDEPVEQNFVRKHALEQAEALGIDIREA        1138
Nbe.v1.1.chr04g34700      RVDVVVNCSGVFRDLFINQMNLLDRAVKMVAELDEPEDQNYVRKHALEQAKTLGVDVREA        1139
Nbe.v1.1.chr14g15590      RVDVVVNCSGVFRDLFINQMNLLDRAVKMVAELDEPEDQNYVRKHALEQAKTLGVDVREA        1139
                          *:***********************:********* :**:*********::**:*:***

AtGUN5_AT5G13630          ATRVFSNASGSYSANISLAVENSSWNDEKQLQDMYLSRKSFAFDSDAPGAGMAEKKQVFE        1198
Nbe.v1.1.chr04g34700      ATRIFSNASGSYSSNINLAVENSTWNDEKQLQDMYLSRKSFAFDCDAPGTGMTEKRKVFE        1199
Nbe.v1.1.chr14g15590      ATRIFSNASGSYSSNINLAVENSTWNDEKQLQDMYLSRKSFAFDCDAPGVGMTEKRKVFE        1199
                          ***:*********:**.******:********************.****.**:**::***

AtGUN5_AT5G13630          MALSTAEVTFQNLDSSEISLTDVSHYFDSDPTNLVQSLRKDKKKPSSYIADTTTANAQVR        1258
Nbe.v1.1.chr04g34700      MALSTADATFQNLDSSEISLTDVSHYFDSDPTNLVQNLRKDGKKPSAYIADTTTANAQVR        1259
Nbe.v1.1.chr14g15590      MALSTADATFQNLDSSEISLTDVSHYFDSDPTNLVQNLRKDGKKPSAYIADTTTANAQVR        1259
                          ******:.****************************.**** ****:*************

AtGUN5_AT5G13630          TLSETVRLDARTKLLNPKWYEGMMSSGYEGVREIEKRLSNTVGWSATSGQVDNWVYEEAN        1318
Nbe.v1.1.chr04g34700      TLSETVRLDARTKLLNPKWYEGMLSTGYEGVREIEKRLTNTVGWSATSGQVDNWVYEEAN        1319
Nbe.v1.1.chr14g15590      TLSETVRLDARTKLLNPKWYEGMLSTGYEGVREIEKRLTNTVGWSATSGQVDNWVYEEAN        1319
                          ***********************:*:************:*********************

AtGUN5_AT5G13630          STFIQDEEMLNRLMNTNPNSFRKMLQTFLEANGRGYWDTSAENIEKLKELYSQVEDKIEG        1378
Nbe.v1.1.chr04g34700      TTFIQDEEMLNRLMNTNPNSFRKLLQTFLEANGRGYWETSAENIEKLKQLYSEVEDKIEG        1379
Nbe.v1.1.chr14g15590      TTFIQDLEMLNRLMNTNPNSFRKLLQTFLEANGRGYWETSEENIEKLKQLYSEVEDKIEG        1379
                          :***** ****************:*************:** *******:***:*******

AtGUN5_AT5G13630          IDR  1381
Nbe.v1.1.chr04g34700      IDR  1382
Nbe.v1.1.chr14g15590      IDR  1382
                          ***
```

**Figure S2**. Multiple sequence alignment of *N. benthamiana* GUN5 homeologous proteins with *A. thaliana* GUN5 (AtGUN5). Nbe.v1.1.chr04g34700 and Nbe.v1.1.chr14g15590 were retrieved from the NbenBase DB [1] using the AtGUN5 sequence as a query. The conserved GUN5 domain was identified by the NCBI Conserved Domain Database (CDD) [2]. GUN5 domain is highlighted with gray for AtGUN5, yellow for Nbe.v1.1.chr04g34700, and blue for Nbe.v1.1.chr14g15590.

Supporting Figure S3

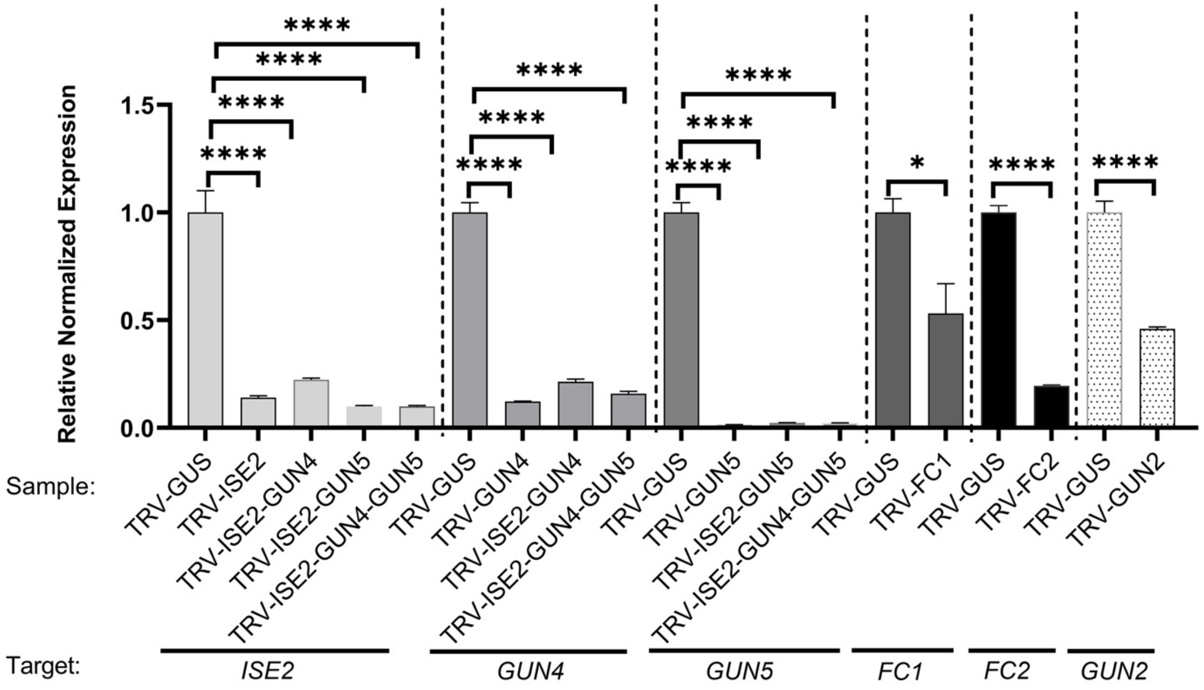


**Figure S3. Confirmation of VIGS.** The relative level of expression of genes targeted for silencing by VIGS was measured by qPCR. Statistical significance was calculated by Student's t-test compared to the non-silenced control (TRV-*GUS*). *= p<0.05, ****p<0.001. Primers used are described in **S1 Table**. Error bars indicate standard deviation, and the results are from three biological replicates.

Supporting Figure S4

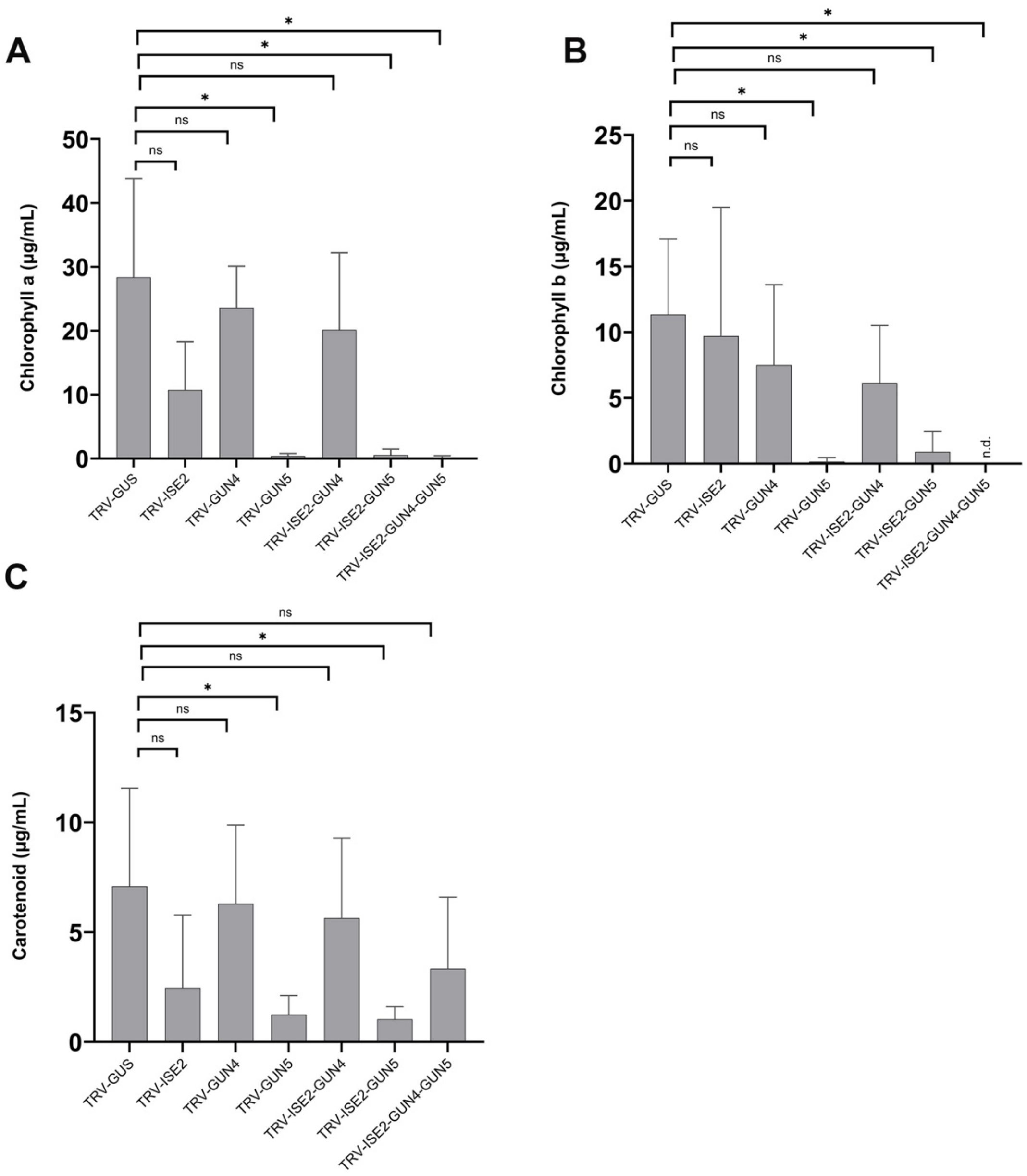


**Figure S4. Chlorophyll a and b, and total carotenoid content of silenced plants.** (a) Chlorophyll a, and (b) chlorophyll b were measured in leaves of silenced plants. (c) Total carotenoid content of leaves of silenced plants. Asterisk denotes statistical significance, Student's t -test ($p<0.05$). ns= not significant.

Supporting Figure S5

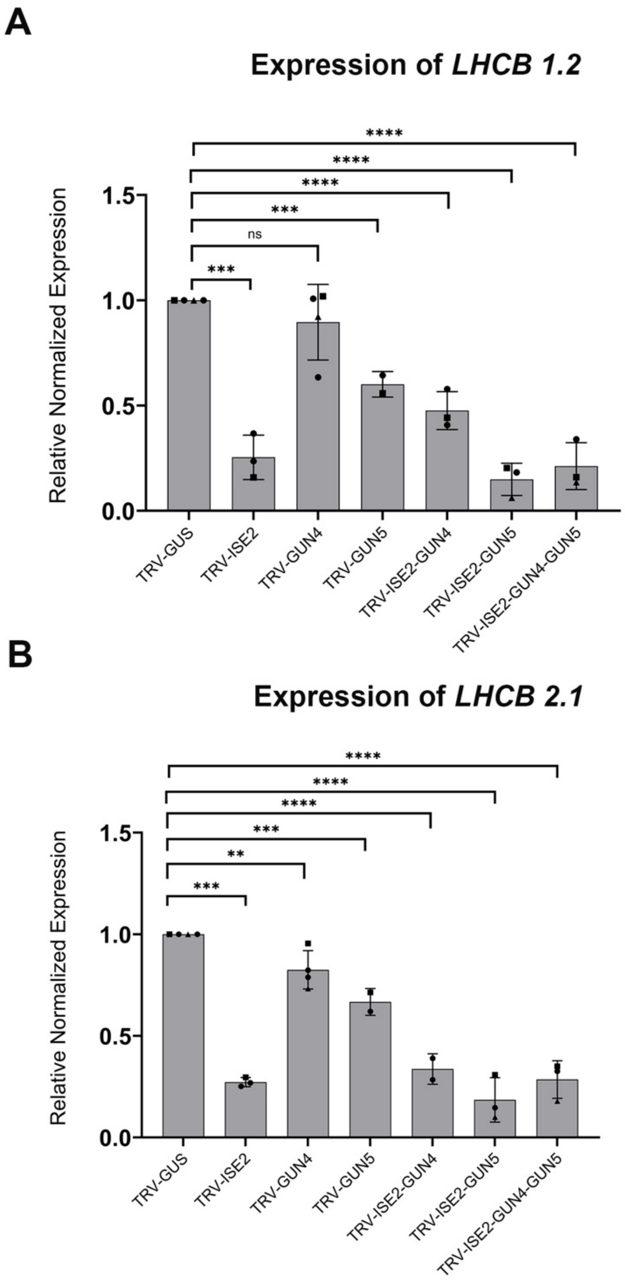


**Figure S5. PhANG expression in silenced plants.** Expression of (a) *Lhcb1.2* and (b) *Lhcb2.1* was measured in silenced plants by qPCR. Statistical significance was calculated by Student's t-test compared to the non-silenced control (TRV-*GUS*). ** indicates $p<0.01$, *** indicates $p<0.001$ and **** indicates $p<0.0001$. Primers used are described in **S1 Table**. Error bars indicate standard deviation, and the results are from three biological replicates.

Supporting Figure S6

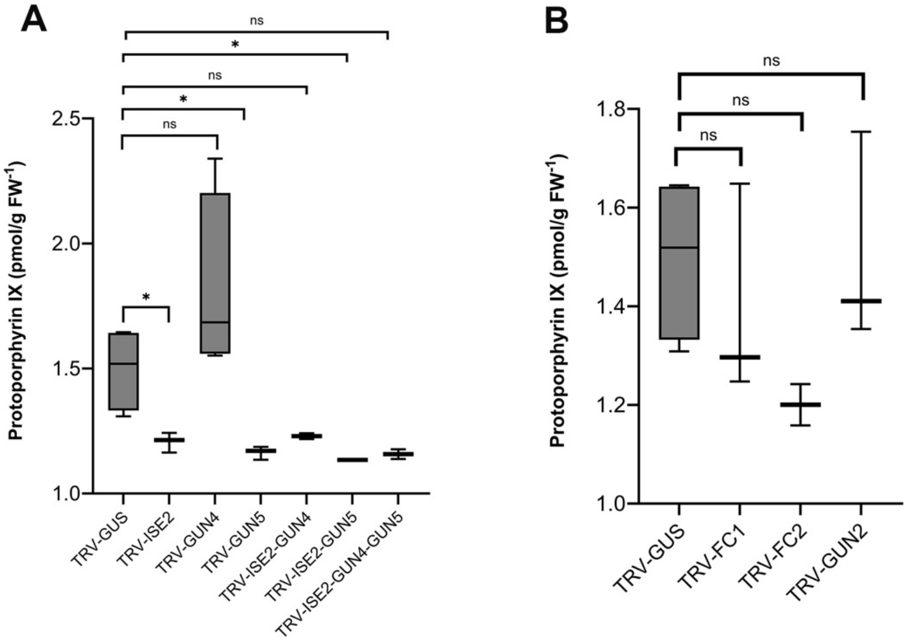


**Figure S6. Protoporphyrin IX measurement in silenced plants.** Proto IX was measured in silenced plants. Data shown is the mean of three biological replicates each consisting of a pool of three plants. Error bars indicate standard deviation. Statistical significance was determined by Student's t test and comparison was made to the non-silenced control (TRV-*GUS*). * indicates $p<0.05$. ns = not significant.

Supporting Figure S7

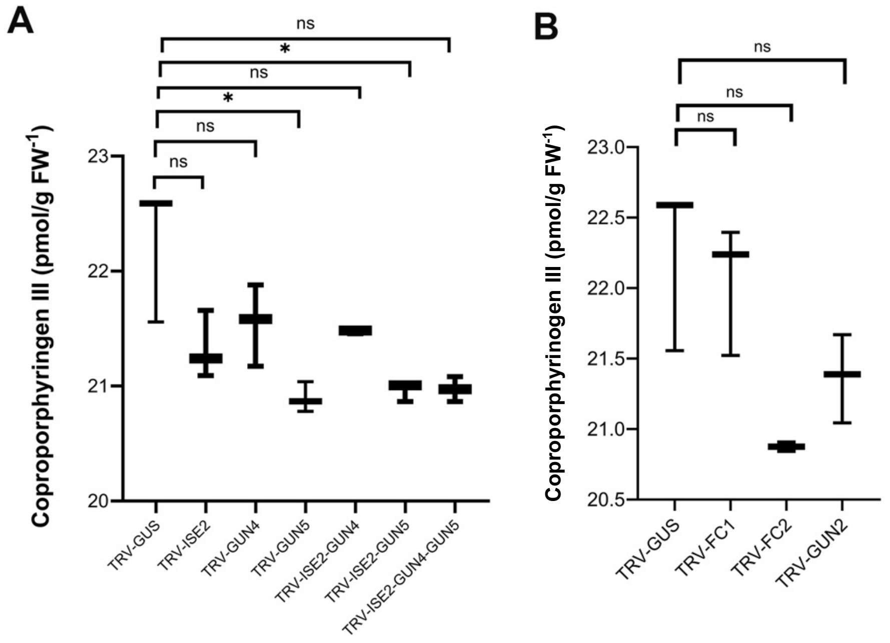


**Figure S7. Coproporphyrinogen III measurement in silenced plants.** Copro III was measured in silenced plants. Data shown is the mean of three biological replicates each consisting of a pool of three plants. Error bars indicate standard deviation. Statistical significance was determined by Student's t test and comparison was made to the non-silenced control (TRV-*GUS*). * indicates p<0.05. ns = not significant.

Supporting Figure S8

```
AtFC1_AT5G26030           MQATALSSGFNPLTKRKDH--R----FPRSCSQRN---------------------SLSL    33
Nbe.v1.1.chr10g28700      MEAGTISRLLQPVKFQSSNLSNFSSNSSMFCPQTRSMPACHASHGGRETNPTEQELSLVL    60
Nbe.v1.1.chr17g40820      MEAGTISRLLQPMKQ----TSNFSSNSSMFCPQTRLMPACHVSHGGRETNPIEQELSMVL    56
                          *:* ::*  ::*:.       .        * * .                     *: *

AtFC1_AT5G26030           IQCDIKERSFGESMTITNRGLSFKTNVFEQAR-----SVTGDCSYDETSAKARSHVVAED    88
Nbe.v1.1.chr10g28700      SSENMKSSVF--------GGLSLHHPVHKRDPVGKMFCSVGSYTYPGSIA-ESPSQTTEE    111
Nbe.v1.1.chr17g40820      SSESKKSSVF--------GGLSLHRPVHKRGPVSKTFCSVGAYTYPGSIA-ESPSQTTEE    107
                           . . *.  *         ***::  *.::       . .*  :*  : *      .:*:

AtFC1_AT5G26030           KIGVLLLNLGGPETLNDVQPFLYNLFADPDIIRLPRPFQFLQGTIAKFISVVRAPKSKEG    148
Nbe.v1.1.chr10g28700      KIGVLLLNLGGPDTLHDVQPFLFNLFADPDIIRLPRLFRFLQRPLAQLISVLRAPKSKEG    171
Nbe.v1.1.chr17g40820      KIGVLLLNLGGPDTLHDVQPFLFNLFADPDIIRLPRLFRFLQRPLAQLISVLRAPKSKEG    167
                          ************:**:******:************* *:***  :*::***:********

AtFC1_AT5G26030           YAAIGGGSPLRKITDEQADAIKMSLQAKNIAANVYVGMRYWYPFTEEAVQQIKKDKITRL    208
Nbe.v1.1.chr10g28700      YAAIGGGSPLRKITDEQASALKMALETKEVPSNVYVAMRYWHPFTEEAVHQIKRDGITKL    231
Nbe.v1.1.chr17g40820      YAAIGGGSPLRKITDEQASALKMALETKEVPANVYVAMRYWHPFTEEAVHQIKRDGITKL    227
                          ******************.*:**:*::*:: :****.****:*******:***:* **:*

AtFC1_AT5G26030           VVLPLYPQYSISTTGSSIRVLQDLFRKDPYLAGVPVAIIKSWYQRRGYVNSMADLIEKEL    268
Nbe.v1.1.chr10g28700      VVLPLYPQYSISTTGSSVRALQNIFKDDSYLSRLPVAIIESWYQRQGYIKSMADLIEKEL    291
Nbe.v1.1.chr17g40820      VVLPLYPQYSISTTGSSVRALQNIFKGDSYLSRLPVAIIESWYQRQGYIKSMADLIENEL    287
                          *****************:*.**::*: * **: :*****:*****:**::*******:**

AtFC1_AT5G26030           QTFSDPKEVMIFFSAHGVPVSYVENAGDPYQKQMEECIDLIMEELKARGVLNDHKLAYQS    328
Nbe.v1.1.chr10g28700      HNFSNPEEVMVFFSAHGVPVSYVEDAGDPYRDQMEECISLIMNELKARGTNNDHTLAYQS    351
Nbe.v1.1.chr17g40820      HNFSNPEEVVVFFSAHGVPVSYVEDAGDPYRDQMEECISLIMNELKARGTNNDHTLAYQS    347
                          :.**:*:**::*************:*****:.******.***:******. ***.*****

AtFC1_AT5G26030           RVGPVQWLKPYTDEVLVDLGKSGVKSLLAVPVSFVSEHIETLEEIDMEYRELALESGVEN    388
Nbe.v1.1.chr10g28700      RVGPVQWLKPYTDEVLVELGKKGVKSLLAVPVSFVSEHIETLEEIDMEYKELALESGIEK    411
Nbe.v1.1.chr17g40820      RVGPVQWLKPYTDEVFVELGKKGVKSLLAVPVSFVSEHIETLEEIDMEYKELALESGIEN    407
                          ***************:*:***.***************************:*******:*:

AtFC1_AT5G26030           WGRVPALGLTPSFITDLADAVIESLPSAEAMSNPNAVVDSEDSESSDAFSYIVKMFFGSI    448
Nbe.v1.1.chr10g28700      WGRVPALNCNSSFITDLADAVIEALPSAMAMSTSTST---EEEV-DDPMQYFIKMLFGSL    467
Nbe.v1.1.chr17g40820      WGRVPALNCTSSFITDLADAVIEVLPSAMAMSTSSGA---EEEVDNDPMQYFIKMLVGSL    464
                          *******. . ************ **** ***. ...   *:.  .* :.*::**:.**:

AtFC1_AT5G26030           LAFVLLLSPKMFHAFRNL--      466
Nbe.v1.1.chr10g28700      LAFVFLFSPKMVSAFRKNIL      487
Nbe.v1.1.chr17g40820      LAFVFLFSPKMVSAFRKNIL      484
                          ****:*:****. ***:
```

**Figure S8**. Multiple sequence alignment of *N. benthamiana* FC1 homeologous proteins with *A. thaliana* FC1 (AtFC1). Nbe.v1.1.chr10g28700 and Nbe.v1.1.chr17g40820 were retrieved from the NbenBase DB [1] using the AtFC1 sequence as a query. The conserved FC1 domain identified by NCBI Conserved Domain Database (CDD) [2]. FC1 domain is highlighted with gray for AtFC1, yellow for Nbe.v1.1.chr10g28700, and blue for Nbe.v1.1.chr17g40820.

Supporting Figure S9

```
AtFC2_AT4G24740          -----MNCPAMTASPSSSSSSSYSTFRPPPPLLPQLSNDSQRSVVMHCTRLPFEAFAATS     55
Nbe.v1.1.chr09g34020     MAPPVIFGREISASPAN--CTSYSPFH-------QVDCQSSVPSAASCLRVPKNCSLVAA     51
Nbe.v1.1.chr11g29350     MAQPVIFCREISASPAN--CTPYSPFH-------QVDGQSSVPFAASCLRVPKNCSLVAA     51
                              :    ::***:.  .: ** *:       *:. :*.   .  * *:* :.  .::

AtFC2_AT4G24740          SNRLLGKHSLPLRAALVTSNPLNISSSSVISDAISSSSVITDDAKIGVLLLNLGGPETLD     115
Nbe.v1.1.chr09g34020     RCSL---KPSMMPSALVTSRPQQVST---------DPVNVVRDGKIGVLLLNLGGPESLE     99
Nbe.v1.1.chr11g29350     RCSL---KPSMMPSALVTSRPQQVST---------NPVNVVCDGKIGVLLLNLGGPESLE     99
                            *   :   : :*****.* ::*:         .    :.*.*************:*:

AtFC2_AT4G24740          DVQPFLFNLFADPDIIRLPPVFQFLQKPLAQFISVARAPKSKEGYASIGGGSPLRHITDA     175
Nbe.v1.1.chr09g34020     DVQPFLFNLFADPDIIRLPRLFRFLQRPLAQFISVARAPKSKEGYASIGGGSPLRRITDA     159
Nbe.v1.1.chr11g29350     DVQPFLFNLFADPDIIRLPRLFRFLQRPLAQFISVARAPKSKEGYASIGGGSPLRRITDA     159
                         ******************* :*:***:****************************:****

AtFC2_AT4G24740          QAEELRKCLWEKNVPAKVYVGMRYWHPFTEEAIEQIKRDGITKLVVLPLYPQFSISTSGS     235
Nbe.v1.1.chr09g34020     QAEALRKALWERNVPAKVYVGMRYWHPFTEEAIELIKRDGITKLVVLPLYPQFSISTSGS     219
Nbe.v1.1.chr11g29350     QAEALRKALWERNVPAKVYVGMRYWHPFTEEAIELIKRDGITKLVVLPLYPQFSISTSGS     219
                         *** ***.***:********************** *************************

AtFC2_AT4G24740          SLRLLERIFREDEYLVNMQHTVIPSWYQREGYIKAMANLIQSELGKFGSPNQTCFKPKYM     295
Nbe.v1.1.chr09g34020     SLRLLESIFREDEYLVNMQHTVIPSWYQREGYIKAMADLMEKELKSFERPGE--------     271
Nbe.v1.1.chr11g29350     SLRLLESIFREDEYLVNMQHTVIPSWYQREGYIKAMADLMEKELKSFEHPGE--------     271
                         ****** ******************************:*::.** .*  *.:

AtFC2_AT4G24740          LQVVIFFSAHGVPLAYVEEAGDPYKAEMEECVDLIMEELDKRKITNAYTLAYQSRVGPVE     355
Nbe.v1.1.chr09g34020     --VMIFFSAHGVPLAYVEEAGDPYKAEMEECVDLIMEELEKRRVHNAYTLAYQSRVGPVE     329
Nbe.v1.1.chr11g29350     --VMIFFSAHGVPLAYVEEAGDPYKAEMEECVDLIMEELEKRRVHNAYTLAYQSRVGPVE     329
                           *:***********************************:**:: ***************

AtFC2_AT4G24740          WLKPYTEEAITELGKKGVENLLAVPISFVSEHIETLEEIDVEYKELALKSGIKNWGRVPA     415
Nbe.v1.1.chr09g34020     WLKPYTDETIIELGKKGVKSILAVPISFVSEHIETLEEIDVEYKELALESGIQNWARVPA     389
Nbe.v1.1.chr11g29350     WLKPYTDETIIELGKKGVKSILAVPISFVSEHIETLEEIDVEYKELALESGIQNWARVPA     389
                         ******:*:* *******:.:***************************:***:**.****

AtFC2_AT4G24740          LGTEPMFISDLADAVVESLPYVGAMAVSNLEARQSLVPLGSVEELLATYDSQRRELPAPV     475
Nbe.v1.1.chr09g34020     LGVEPTFISDLADAVIESLPYVGAMAVSNLEARQSLVPLGSVEELLAVYDSQRRELPPPV     449
Nbe.v1.1.chr11g29350     LGVEPTFISDLADAVIESLPYVGAMAVSNLEARQSLVPLGSVEELLAAYDSQRRELPPPV     449
                         **.** *********:*******************************.********* **

AtFC2_AT4G24740          TMWEWGWTKSAETWNGRAAMLAVLALLVLEVTTGKGFLHQWGILPSL-------------     522
Nbe.v1.1.chr09g34020     TVWEWGWTKSAETWNGRAAMLAVLVLLVLEVTTGEGFLHQWGILPLFP------------     497
Nbe.v1.1.chr11g29350     TFWEWGWTKSAETWNGRAAMLAVLVLLVLEVTTGEGFLHQWVFAAKKPSFHSLEASEQVS     509
                         *.**********************.*********:****** :

AtFC2_AT4G24740          -------        522
Nbe.v1.1.chr09g34020     -------        497
Nbe.v1.1.chr11g29350     EIHSTLE        516
```

**Figure S9**. Multiple sequence alignment of *N. benthamiana* FC1 homeologous proteins with *A. thaliana* FC2 (AtFC2). Nbe.v1.1.chr09g34020 and Nbe.v1.1.chr11g29350 were retrieved from the NbenBase DB [1] using AtFC2 sequence as a query. The conserved FC2 domain identified by NCBI Conserved Domain Database (CDD) [2]. The FC2 is highlighted with gray for AtFC2, yellow or Nbe.v1.1.chr09g34020, and blue for Nbe.v1.1.chr11g29350.

Supporting Figure S10

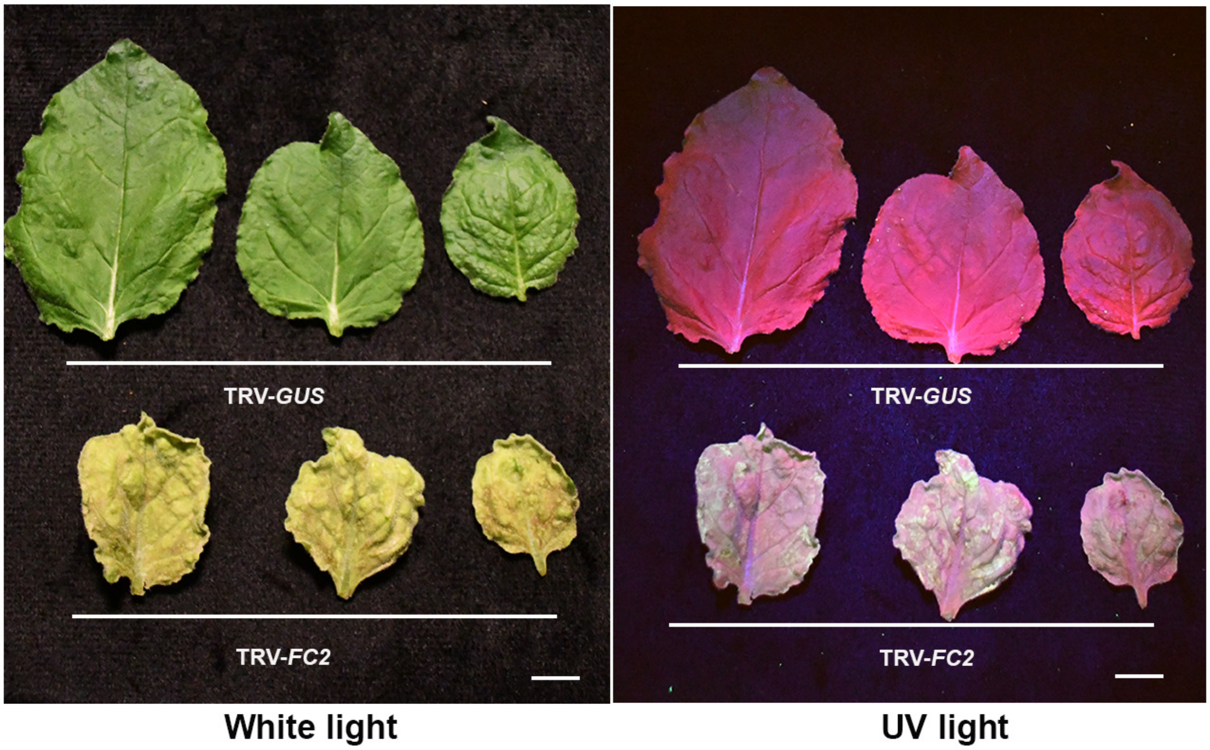


**Figure S10**. **Cell death in *FC2*-silenced plants.** Compared to a non-silenced control plant (TRV-*GUS*), the FC2-silenced plant (TRV-*FC2*) shows autofluorescence under UV light, indicating cell death.

**Table S5. Arabidopsis mutants used in this study**

| Mutant | Background | Reference |
|---|---|---|
| *gun2* | Col-o | Susek et al., 1993 |
| *gun4-1* | Col-0 | Koussevitzky et al., 2007 |
| *gun5-1* | Col-0 | Koussevitzky et al., 2007 |

**Table S6. Primers used in this study**

| Gene | Forward sequence | Reverse sequence | Purpose |
|---|---|---|---|
| *NbGUN2** | gattctagaATGGCTTCAATTACACCCTTATC | gatctcgagTCAAGACAATATTAATCGGAGG | VIGS construct for silencing |
| *NbGUN2** | TTCTCAGGGGAAATCCTGCG | TGAGCTCCAATGTACAACTTACC | qPCR for confirmation of silencing |
| *NbFC1* | ATCggatccGCTAAAAATGGCTCTAGAAAC | TGAgaattcAGAACCAGTTGTAGAGATAG | VIGS construct for silencing |
| *NbFC1* | CGATTTCTCCAACGCCCATT | TTGCAGCGTACCCTTCCTTA | qPCR for confirmation of silencing |
| *NbFC2* | ATCgaattcTGGGAAAGGAATGTGCCTGC | TGAggatccCCACACATTCCTCCATTTCA | VIGS construct for silencing |
| *NbFC2* | GCTCGTCAGTCATTGGTTCC | GCCAACATAGCAGCTCTTCC | qPCR for confirmation of silencing |
| *NbGUN4* | TAAgaattcGGAGAAGGCATTCTATTGATTGCCC | AGCggatccCACTTTGTTCCATATTTTCTTTTG | VIGS construct for silencing |
| *NbGUN4* | ACAAGCAAGTTTTTCTGGGGA | ACTCCTCAAACCACCCAACA | qPCR for confirmation of silencing |
| *NbGUN5* | ATGgaattcGCTCAGCGAGCTCATTTCCTC | ATCggatccGGTTAGCTTGTCATTGACATTCAC | VIGS construct for silencing |
| *NbGUN5* | CCGAGCAGTCAAGATGGTTG | GCTTGTTCTAGTGCATGTTTCC | qPCR for confirmation of silencing |
| *NbLHCB1.2*** | CAGATGGGCTATGCTTGGTG | GCCTTGAACCATACAGCCTC | qPCR primer |
| *NbLHCB2.1*** | AGGCCCACTTGGTGAAGG | TCGGGATCATCAGCGAAC | qPCR primer |
| *NbEF1α*** | CAAGAGGCCCTCAGACAAAC | CACAACCATACCAGGCTTGA | qPCR primer |
| *NbRH22*** | CTAtctagaGAGCAAGTGGTTCGAATG | CTAggatccGATAACTGCTTCTCATC | VIGS construct for silencing |
| *NbRH39*** | TATtctagaCACCTGAACAATAGGCAACAT | TATggatccAACAGGAAGAGCACTGCTAAA | VIGS construct for silencing |

| | | | |
|---|---|---|---|
| *NbISE2*** | GACctcgagGGGGAAGGTTGATACTTTGA | GATggatccTCAAACGGGAGACCTTATTC | VIGS construct for silencing |
| *NbPNPaseA*** | CGTgaattcATGCCTTGGCAGTTACGGC | CGAtctagaCGACTAGCTTATACAGCTC | VIGS construct for silencing |
| *NbPNPaseA/B*** | CGTgaattcGATGCATCAGGAGATATGG | CGAtctagaCGTCCAGGCGCAATCTCCA | VIGS construct for silencing |
| *NbRNaseJ*** | GTAtctagaGGCCTTGAGGAACTGTC | CATggatccCAGAGTTGATGGATCTATTG | VIGS construct for silencing |
| *NbIPI1*** | CACgaattcCCTGTGTCCTTGTGATATC | CATtctagaCATAAGAAGCCCAAAGGTTA | VIGS construct for silencing |
| *NbRH3*** | AAATTTggatccTTTTGTTAGGGCATTCGGATG | CTTAAtctagaAAAATTCACGTTCAGGGTGCT | VIGS construct for silencing |
| *NbPDS*** | CGAggatccCTGACGAGCTTTCGATGCAGTGCAT | TCGctcgagATATATGGACATTTATCACAGGAAC | VIGS construct for silencing |

* From [3]

** From [4]

**Table S7. Plasmids used in this study**

| Plasmid | Description |
|---|---|
| pTBS13* | *ISE2* silencing construct in pTRV1 (pYL156) for VIGS |
| pLBG269 | *GUN4* silencing construct in pTRV1 (pYL156) for VIGS |
| pLBG270 | *GUN5* silencing constructs in pTRV1 (pYL156) for VIGS |
| pMFA337 | *ISE2* and *GUN4* silencing constructs in pTBS13 for VIGS |
| pMFA338 | *ISE2* and *GUN5* silencing constructs in pTBS13 for VIGS |
| pMFA340 | *ISE2* and *GUN4* and *GUN5* silencing constructs in pTBS13 for VIGS |
| pMFA342 | *FC1* silencing constructs in pTRV1 (pYL156) for VIGS |
| pMFA343 | *FC2* silencing constructs in pTRV1 (pYL156) for VIGS |
| pMFA345 | *GUN2* silencing constructs in pTRV1 (pYL156) for VIGS |

Supporting Methods

**Chlorophyll and carotenoid determination**

Chlorophyll and carotenoid were extracted following the method described previously [4]. Briefly, ten leaf discs were weighed and homogenized in 3 mL of 100% acetone using a mortar and pestle. Two mL of the resulting extracts were transferred to a reaction tube and centrifuged at 500 x g for 5 minutes. The supernatants were used to measure absorption at 470 nm, 646.8 nm, and 661.6 nm. Quantification of total carotenoids (x+c, xanthophylls, and carotenes) and chlorophyll a and b were performed using the equation described in reference [5].

**Porphyrin measurements**

Protoporphyrin IX (Proto) and coproporphyrinogen III (copro III) were measured by HPLC analysis as described in [6]. Briefly, ten leaf discs from each VIGS plant were weighed to measure the fresh weight and then frozen in liquid nitrogen. The discs were ground using mortar and pestle, and homogenized were resuspended with 250 µL of 50 mM potassium phosphate buffer, pH 7.8, and centrifuged at 10 000 x g for 10 minutes at 4'C. The supernatant was transferred to a dark tube and kept on ice. To extract tetrapyrrole from the pellet, an additional 500 µL 90% (v/v) methanol and 10% (v/v) 0.1 M NH4OH were added, followed by centrifugation at 10, 000 x g for 10 minutes at 4'C. The supernatants were combined, and tetrapyrroles were extracted from the pellet again with 500 µL 90% (v/v) acetone and 10% (v/v) 0.1 M $NH_4OH$. This mixture was incubated at 20'C for 20 minutes and then centrifuged at 16,000 x g for 10 minutes at 4'C. All the extracts were combined and subjected to final centrifugation at 16,000 x g for 10 minutes at 4'C. To convert copro III to fluorescent (detectable) porphyrins, the 400 µL extracts were mixed with 10 µL 1M acetic acid and 10 µL of 2-butanone for oxidation prior to injection into HPLC.

Tetrapyrroles were separated using a Waters Symmetry C18 column (5 µm, 4.6 x 150 mm) with the following HPLC method: 7 min at 100% Solvent A (10% (v/v) methanol and 10% (v/v) 1M ammonium acetate, pH 5.2), 13 min 100% Solvent B (90% (v/v) methanol and 10% (v/v) 1 M ammonium acetate, pH 5.2), 9 min 100% Solvent A. Fluorescence detection was performed at an excitation/emission wavelength of 405/625 nm for copro III and Proto IX. Standard curves were generated using authentic tetrapyrroles (Frontier Scientific, Logan, UT).

**Gene Expression Analysis**

Gene ontology (GO) and KEGG pathway enrichment analyses of DEGs were conducted using KOBAS website (http://kobas.cbi.pku.edu.cn) with Arabidopsis genomes [7]. GO enrichment analyses utilized the hypergeometric statistical model, and p-values were adjusted using the Benjamini-Hochberg method to control the false discovery rate (FDR). GO terms and KEGG pathways with FDR < 0.05 were considered significantly enriched.

**Supplemental References**


1. Kurotani KI, Hirakawa H, Shirasawa K, Tanizawa Y, Nakamura Y, Isobe S, et al. Genome Sequence and Analysis of Nicotiana benthamiana, the Model Plant for Interactions between Organisms. Plant Cell Physiol. 2023;64(2):248-57. doi: 10.1093/pcp/pcac168. PubMed PMID: 36755428; PubMed Central PMCID: PMCPMC9977260.

2. Wang J, Chitsaz F, Derbyshire MK, Gonzales NR, Gwadz M, Lu S, et al. The conserved domain database in 2023. Nucleic acids research. 2023;51(D1):D384-D8. doi: 10.1093/nar/gkac1096. PubMed PMID: 36477806; PubMed Central PMCID: PMCPMC9825596.

3. Brunkard JO, Runkel AM, Zambryski PC. Chloroplasts extend stromules independently and in response to internal redox signals. Proc Natl Acad Sci U S A. 2015;112(32):10044-9. doi: 10.1073/pnas.1511570112. PubMed PMID: 26150490; PubMed Central PMCID: PMC4538653.

4. Ganusova EE, Reagan BC, Fernandez JC, Azim MF, Sankoh AF, Freeman KM, et al. Chloroplast-to-nucleus retrograde signalling controls intercellular trafficking via plasmodesmata formation. Philos Trans R Soc Lond B Biol Sci. 2020;375(1801):20190408. Epub 2020/05/05. doi: 10.1098/rstb.2019.0408. PubMed PMID: 32362251; PubMed Central PMCID: PMCPMC7209952.

5. Lichtenthaler HK, Wellburn AR. Determination of total carotenoids and chlorophylls *a* and *b* of leaf extracts in different solvents. Biochemical Society transactions. 1983;11:591-2. doi: 10.1042/bst0110591.

6. Czarnecki O, Peter E, Grimm B. Methods for analysis of photosynthetic pigments and steady-state levels of intermediates of tetrapyrrole biosynthesis. Methods Mol Biol. 2011;775:357-85. doi: 10.1007/978-1-61779-237-3_20. PubMed PMID: 21863454.

7. Bu D, Luo H, Huo P, Wang Z, Zhang S, He Z, et al. KOBAS-i: intelligent prioritization and exploratory visualization of biological functions for gene enrichment analysis. Nucleic acids research. 2021;49(W1):W317-W25. doi: 10.1093/nar/gkab447. PubMed PMID: 34086934; PubMed Central PMCID: PMCPMC8265193.
